# Evaluation of network architecture and data augmentation methods for deep learning in chemogenomics

**DOI:** 10.1101/662098

**Authors:** Benoit Playe, Véronique Stoven

## Abstract

Among virtual screening methods that have been developed to facilitate the drug discovery process, chemogenomics presents the particularity to tackle the question of predicting ligands for proteins, at at scales both in the protein and chemical spaces. Therefore, in addition to to predict drug candidates for a given therapeutic protein target, like more classical ligand-based or receptor-based methods do, chemogenomics can also predict off-targets at the proteome level, and therefore, identify potential side-effects or drug repositioning opportunities. In this study, we study and compare machine-learning and deep learning approaches for chemogenomics, that are applicable to screen large sets of compounds against large sets of druggable proteins. State-of-the-art drug chemogenomics methods rely on expert-based chemical and protein descriptors or similarity measures. The recent development of deep learning approaches enabled to design algorithms that learn numerical abstract representations of molecular graphs and protein sequences in an end-to-end fashion, i.e., so that the learnt features optimise the objective function of the drug-target interaction prediction task. In this paper, we address drug-target interaction prediction at the druggable proteome-level, with what we define as the chemogenomic neuron network. This network consists of a feed-forward neuron network taking as input the combination of molecular and protein representations learnt by molecular graph and protein sequence encoders. We first propose a standard formulation of this chemogenomic neuron network. Then, we compare the performances of the standard chemogenomic network to reference deep learning or shallow (machine-learning without deep learning) methods. In particular, we show that such a representation learning approach is competitive with state-of-the-art chemogenomics with shallow methods, but not ultimately superior. We evaluate the most promising neuron network architectures and data augmentation techniques, such as multi-view and transfer learning, to improve the prediction performance of the chemogenomic network. Our results shed new insights on the design of chemogenomics approaches based on representation learning algorithms. Most importantly, we conclude from our observations that a promising research direction is to integrate heterogeneous sources of data such as various bioactivity datasets, or independently, multiple molecule and protein attribute views, instead of focusing on sophisticated, yet intuitively relevant, encoder’s neuron network architecture.

## 1 Introduction

The current paradigm in rationalised drug design is to associate a disease to one or several molecular targets, usually proteins called target proteins, or therapeutic targets, involved in biological pathways necessary to the disease development. The goal of the drug discovery process is then to identify a small molecular compound that binds the target protein in such a way that it alters the disease development. This small molecule must also be able to reach the protein target through the organism without triggering undesired deleterious side-effects.

Despite all the advances in understanding diseases, and technological breakthroughs, this rational drug discovery process has limited success, and only a few tens of new drugs reach the market every year. Single-drug development is currently estimated to require about 1.8 billion US dollars over about 13 years on average [1]. The main bottleneck of the drug design process is the toxicity issue. Most of the hits identified by high-throughput screening (HTS) fail to become approved drugs because of unexpected side effects and toxicity, which are in part due to unexpected interactions of the drug with so called “off-target” proteins. Moreover, while HTS helps increase the number of tested molecules to identify hits, it is not possible to conduct bio-assays at the human proteome scale [2, 3]. In addition, more than 6 million drug-like compounds are commercially available, but the drug-like molecule space is estimated to be around 10^60^ [4]. Randomly exploring this space is out of reach even by HTS, and identifying a molecule in such a vast space relies mainly on human expertise.

In that context, computational approaches appear as promising ways to facilitate the tedious task of drug discovery. In particular, chemoinformatics [5] algorithms provide in silico screening methods that help and guide the drug discovery process.

Such in silico drug virtual screening approaches can help identify compounds that would bind to the considered therapeutic target to guide experiments and only test potentially active combinations, thus reducing the cost and time required for HTS. Such methods are called ligand-based drug virtual screening approaches, or Quantitative Structure-Activity Relationship (QSAR) methods [6]. They use a set of compounds for which the activity has been experimentally measured and learn a model function to relate the structure of molecules to their biological activities.

Chemogenomics is a valuable generalisation of ligand-based approach where the prediction models for several targets are learned simultaneously [7]. The underlying idea is that the prediction of the drug biological activity is expected to benefit from the activities known for other molecules and other targets. Moreover, chemogenomics can virtually screen drugs at the druggable proteome level, which enables to detect unexpected “off-target” for drugs, which can be related to unwanted side-effects. However, these off-target interaction can sometimes be beneficial in other clinical situations, and suggest new therapeutic indications for drugs. In a nutshell, chemogenomics, as a druggable proteome-wide drug virtual screening approach, can help to identify drug candidates for known therapeutic targets, and is expected to help anticipate the toxicity issues of candidate drugs by detecting side effects-related off-targets. It can also lead to repurposing of marketed drugs in new therapeutic indications.

Therefore, in this study, we focus on chemogenomics with deep learning approaches, for virtual screening of large sets of compounds against large sets of druggable proteins. We formulate the problem as a Drug-Target Interaction (DTI) prediction task, or equivalently drugs “virtual screening” (VS), in terms of a binary classification of protein/molecule pairs that interact or do not interact.

The paper is organised as follows. Section 2 recalls key reference machine-learning methods not involving deep-learning, in the context of chemogenomics. Section 3 presents what we define as our “standard” chemogenomic network. More precisely, we present the different blocks (molecule encoder, protein encoder, prediction feed-forward network) that constitute our deep learning approach followed throughout the paper. In particular, we also introduce the required background and propose a general formulation of recent representation learning-based models which learn descriptors on the molecular graph and protein sequence in an end-to-end fashion. Section 4 shortly reviews related works published until now in chemoinformatics or chemogenomics with deep learning approaches. Sections 5 presents the Materials and Methods.

Section 6 presents our Results. First, using two DrugBank-based datasets, we build over previous works in chemogenomics with deep learning by a more exhaustive evaluation of variants in the chemogenomic network architecture (namely, of the molecule, protein encoders, and of the method that combines both representations), which is a first contribution. In particular, we evaluate the performances of the considered methods in four realistic scenarios, i.e. the general case, but also the protein, molecule, and double orphan cases, in order to better define which method performs better, depending on the considered data. Importantly, we compare the prediction performances to state-of-the-art machine-learning methods that use deep learning, or not, which was not performed in previous studies in chemogenomics with deep learning.

In addition, we also test data augmentation techniques, namely multi-view learning and transfer learning, to enhance representation learning for chemogenomics, which is a new contribution in this field. In particular, we explore the simultaneous use of expert-based and learnt features in a differentiable fashion, with the idea that combining both types of features might increase performances. We also explore transfer learning via curriculum learning. This refers to using an additional larger dataset for pre-training the neuron network with an auxiliary task, before using it in the chemogenomic task of interest, with the aim of learning richer representations of molecules or proteins. We tested various ways to implement this idea.

Section 7 finally presents a short discussion and conclusion about the proposed methods for chemogenomics with deep learning.

Regarding the implementation, our code is available on GitHub at: *https://github.com/bplaye/NNk_chemo*. The code and data are available at: *http://members.cbio.mines-paristech.fr/~bplaye/NNk_chemo.zip*. Also, we broadly used machine learning shallow algorithms and auxiliary functions (for instance for computing performance scores) implemented in the scikit-learn library [8], a viral and well-maintained free software machine learning library for Python. Also, we used the Tensorflow library [9] and the Keras library [10] to implement deep learning algorithms. To implement *NRLMF*, we used the PyDTI python package [11] available at: *https://github.com/stephenliu0423/PyDTI*. We used the implementation of LAkernel software [12] available at: http://members.cbio.mines-paristech.fr/jvert/software/.

## 2 Pioneering works in machine-learning for chemogenomics

Various chemogenomics methods have been proposed in the last decade [13, 14, 15, 16, 17, 18, 19, 20, 21, 22, 23, 24, 25, 26, 11]. They differ by (i) the descriptors used to encode proteins and ligands or the similarities between these objects, (ii) the ML algorithm that is used to learn the model and make the predictions. We shortly describe the principle of those the methods displaying state-of-the-art performances.

Most of them are similarity-based approaches, meaning that they consider as input data the similarity between instances. They rely on the assumption that similar compounds (resp. protein), regarding structure, topology or physical properties etc., have similar functions and bio-activities and, therefore, have similar targets (resp. ligands). This assumption can however be wrong in the context of virtual screening, because a tiny change in the molecular structure or protein sequence can sometimes lead to complete loss of interaction.

Jacob and Vert [7] first used the powerful kernel framework to provide efficient ways to combine protein and ligand representations into a joint chemical and biological space. More precisely, they used the Kronecker product of individual kernels to define the kernel associated with the chemogenomic space (or protein-molecule pairwise space). Such an approach have been successfully applied to DTI prediction [14, 27, 28]. We refer to this method as “kronSVM” in the following.

Matrix factorisation (MF) approaches decompose the interaction matrix that lives in the (protein, molecule) space into the product of matrices of lower ranks, living in the two latent spaces of proteins and molecules. First, Gonen et al. [26, 29] proposed probabilistic matrix factorisation in which matrices are defined as random variables, which leads to a probabilistic inference. Second, Liu et al. developed Neighbour Regularised Logistic Matrix Factorisation [11] (*NRLMF*), which consider Logistic Matrix Factorisation [30]. A specific feature of the *NRLMF* method is that it integrates a neighbourhood regularisation method, which allows taking into account only the nearest neighbours to predict a given (protein, ligand) interaction. *NRLMF* displays excellent performances and is also computationally efficient. Note that the *NRLMF* approach is also generalised to orphan molecules and proteins, by computing latent representations of orphan molecules and proteins as a weighted sum of the latent representation of their neighbours. Based on the results reported in the original paper [11], we observe that the proposed method *NRLMF* outperforms the other state-of-the-art methods [23, 22, 26, 24] on the Yamanishi benchmark dataset [13]. Therefore, this method is used in this study as reference method to compare the proposed deep learning approaches for chemogenomics.

## 3 Deep learning methods for chemogenomics

In the following, we first present our global framework for deep learning in chemogenomics. Then, we shortly review the methods that can be used in the different building blocks of our framework: molecules and proteins encoding, combination of these encoding, and prediction network.

### 3.1 Chemogenomic neuron network

Our general scheme for chemogenomics with deep learning can be represented as shown in Fig. 1. It contains four main building blocks: (1) a molecule encoder that calculates descriptors for molecules, (2) a protein encoder that calculates descriptors for proteins, (3) a *Comb* block, i.e. an operation or a neuron network module that combines molecule and protein descriptors in order to build a pairwise latent representation for the molecule-protein pair, and (4) the *MLP*_*pair*_ (for Multi Layer Percepton on pairs, also called feed-forward neuron network, FNN) that predicts whether or not the molecule-protein pairs interact. Any protein sequence and molecular graph encoders can be considered.

**Figure 1:**
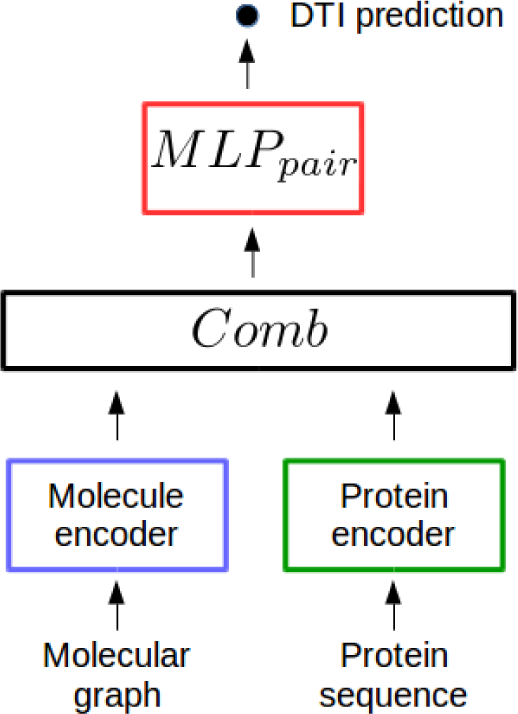
The chemogenomic neuron network.

In the present paper, we refer to this general architecture as a “chemogenomic neuron network”. In the following, we discuss the methods can can be used in these four building blocks.

### 3.2 Molecular graph encoder

Classically, molecules are encoded into a one dimensional representation, like the SMILES [31] representation. However, molecules are also usually represented via their molecular graphs. A molecular graph is an undirected graph *G* = (*𝒱, ε*), where *𝒱* is the set of vertices (or nodes), and *ε* the set of edges. Nodes correspond to atoms and edges to covalent bonds. Thus, molecular graphs belong to the family of attributed undirected graph, since nodes are attributed with atom properties (like atom type category, but also any physicochemical and topological properties), and edges are attributed with bond properties (like bond type, but also topological properties, for instance). Graph Neuron Networks (GNN) are neuron network modules that encode descriptors on undirected graphs, and therefore, they can be used to encode molecular graphs.

Approaches based Recurrent Neuron Network (RNN) modules processing SMILES proved to be competitive with GNN which directly process the molecular graph [32, 33]. However, GNNs appear as a more valuable extension of method repertoire, because of their implementation versatility (as will be explored in the Results section) and because they offer a larger room for improvements. Therefore, in the present study, we used GNN for encoding of molecules.

In the following, we use the word “attribute” to describe the original feature vector of nodes and edges. We use the terms “embedding”, “encoding” or “representation” to refer to the feature vectors of nodes and edges that are learnt by GNNs. We note that each node *i* has an input attribute vector **x**_*i*_ and each edge (*i, j*) has an attribute vector **x**_*ij*_. **h**_*i*_ and **h**_*ij*_ are the node and edge learnt embeddings. 𝒩 (*i*) refers to the neighbouring nodes of node *i*, which can be the one-hop neighbourhood (i.e. the nodes that as reachable by paths of length 1, which we used in practice), and *L* represents the total number of neuron layers in the GNN.

We present our general algorithm for GNN, inspired by Hamilton et al. [34, 35], in Alg. 1.

**Algorithm 1.**
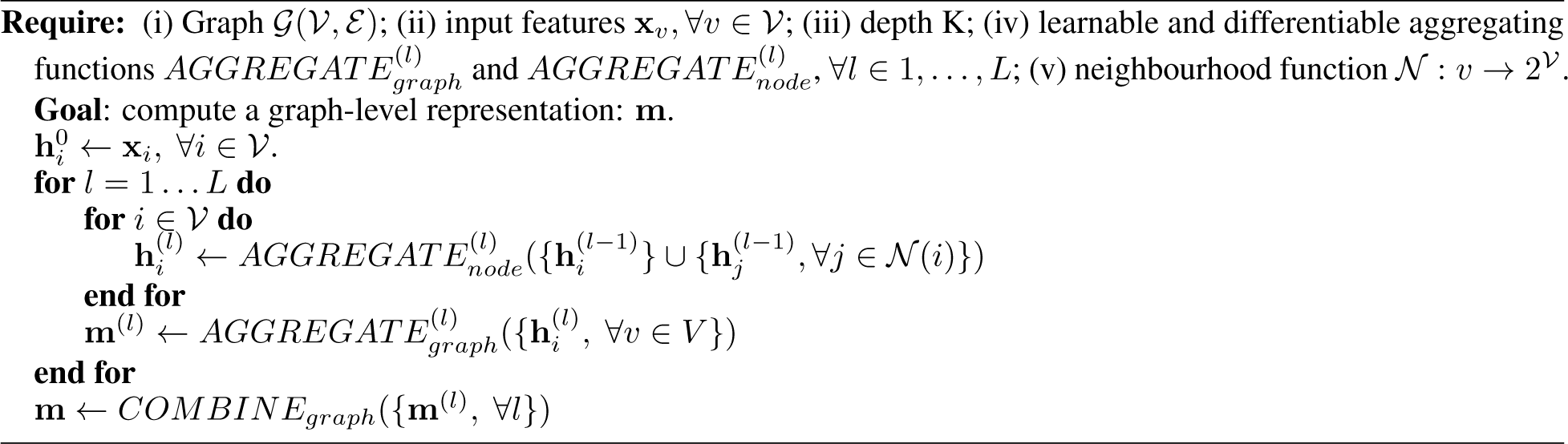
Graph Neuron Network (GNN)

In more explicit words, at each layer *l*, each node aggregates information from its local neighbours in a representation vector 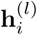 The 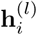 representations for all nodes are aggregated in an **m**^(*l*)^ representation of the molecule. As this process iterates, nodes incrementally gain more information from further parts of the graph, while encoding local graph topology and distribution of attributes. Finally, a global representation of the molecule **m** is derived by combining the **m**^(*l*)^ representations for all considered size of substructures. Various GNN methods can be considered, depending on the aggregation functions that are chosen at the nodes and graph levels, and depending on the function used to combine the **m**^(*l*)^ representations.

Such an approach is spatial and convolutional because it iteratively updates a node representation with a function inputting its neighbourhood, in a manner similar to the receptive field of a convolutional kernel in computer vision.

The learnt representation simultaneously encodes the topological structure of each node’s neighbourhood, as well as the distribution of node features in the neighbourhood.

Note that it is also possible to consider, at each iteration, several neighbourhood sizes by simply duplicating and applying the 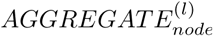 function to several r-hop neighbourhood. The latter is defined as all nodes reachable by paths of length r [36].

Early works [37, 38] showed that graph convolutional networks extract structural features in a similar fashion as standard ECFP procedures [38], and the 1-WL algorithm [37]. Thus, GNNs do not bear new concepts to process graphs, but they learn structural features that optimise the prediction performance.

Other works [39, 40] emphasise that GNNs can, at best, be as discriminative as the 1-WL (in terms of the ability to distinguish non-isomorphic graphs with the extracted features). However, the flexibility of GNNs is expected to enhance their representation power with respect to the problem at hand, and therefore, to enable overtake state-of-the-art performance.

The flexibility and representation power of the GNN general formulation in Alg. 1 relies on the aggregation functions. They update node-level representations based on the nodes in their neighbourhood, and compute a graph-level representation based on all nodes representations.

Let us define what we call the “minimal” formulation of GNN, and introduce succinctly some of the alternatives proposed in the literature.

In this minimal formulation, the most intuitive and simple 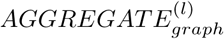 function is used, i.e. the sum over the node representations at all layers [38, 41, 42, 32, 33, 43]:

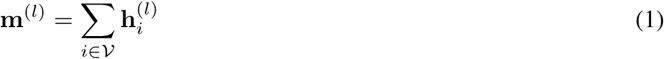

This sum function captures the distribution of each node features in a single value.

In the minimal 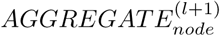 function, the node representation 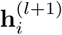 at layer *l* + 1 aggregates representations of its neighbourhood at layer *l* by summing their representations. However, representations of the neighbours are passed through a shared hidden layer before the summation (otherwise, we do not learn any convolutional functions to process the graph). In eq. 2, *σ* refers to the sigmoid function, *𝒩*(*i*) to the one-hop neighbourhood of node *i*, 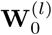 and 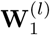 are learnable weight matrices and *α*_*ij*_ are learnable or fixed coefficients.

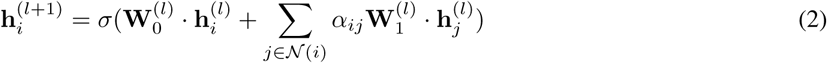

Finally, the minimal *COMBINE*_*graph*_ function used in most studies to provide a graph-level embedding **m** (finally encoding the molecule, in our case), is simply the last layer representation **m**^(*L*)^.

Other 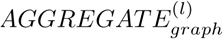 functions than the above minimal function have been proposed in the graph representation learning literature. For instance, another obvious 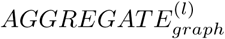 function is the max function, first proposed by [35], which enforce the model to learn molecule-level encodings via the absence or presence of specific key features within the nodes. Alternatively, Li et al. [44] considered a self-attention mechanism to build a graph-level representation, which automatically selects the most relevant nodes for the graph-level task at hand. LSTM-based 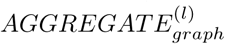 functions have also been proposed [35, 45], which are more flexible than static sum or max functions. Furthermore, Kearnes et al. [46] proposed to design a graph-level description by concatenating the distribution of each feature of the node representations. More precisely, they encoded the distribution of each individual features by a n-bits histogram. From another perspective, several authors [47, 48, 49] proposed to learn a graph-level embeddings by creating a virtual “supernode”, that is connected to all other nodes via a particular type of edge. This virtual node is updated like real nodes, and the final representation of this virtual node is used as the graph representation. In particular, Ishiguro et al. [50] showed that such a supernode can improve the representation power of a wide variety of existing GNNs, by allowing pairs of distant nodes to communicate through the supernode in one hop.

A promising approach to define a suitable 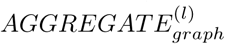 function is based on differential hierarchical pooling, which has been proposed simultaneously by several groups [51, 52, 53]. It allows for the graph convolutional architecture to iteratively operate on coarser representations of a graph. To be precise, such procedure actually replaces both the 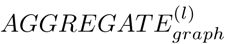 and *COMBINE*_*graph*_ functions, since it results directly in a final graph-level representation. This idea was to design an analogue of the pooling operation for images, which was shown to be of crucial importance for convolutional neuron network success in the field of image processing. Indeed, one of the possible limitations of current GNNs is that they are inherently flat, since they only propagate information across the edges of the graph, without reducing the size of the graph. However, this could be not particularly suitable for molecular graphs. Indeed, they are small enough so that iterative convolutional node updates, followed by a simple summation of graph nodes to build a graph-level embedding, could equivalently encode and hierarchically agglomerate graph substructures of different sizes. Therefore, we discuss and evaluate differential hierarchical pooling in more details later in this work.

Many other 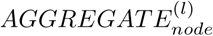 functions have also been proposed in the literature, but they have not been tested in the context of chemoinformatics or chemogenomics. Typically, some studies have considered to sum over edge type-specific standard convolutional layers [53, 54, 55], or even also proposed to update edges attributes similarly to node attributes, hoping for more expressive GNN [56, 57, 53]. We tested this principal, i.e. taking into account bond attributes in the molecule graph, as shown in the Results section.

Velickovic et al. [58] proposed an attention mechanism to design a neighbourhood aggregation function (instead of a node aggregation function) for graph representation learning. Their GAT 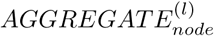function resulted in performance improvements higher than those reported in other studies, which usually show very little performance increase (one or two points maximum) on graph prediction gold standard datasets comparatively to minimal GNN formulation. In addition, this method seems particularly powerful since it aggregates neighbouring information after identifying the most suitable task-specific neighbours at each node. However, we could wonder whether the GAT 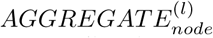 would be interesting in the case of small graphs such as molecular graphs, where all atoms are supposedly relevant. Recently, Shanthamallu et al. [59] quantitatively assessed that the distribution of attention weights for any node obtained by GAT is almost always uniform, in the case of gold standard datasets for graph representation learning. This means that the attention mechanism is not suitable in these cases, since its selects all neighbours as relevant. Therefore, we discuss and evaluate the GAT 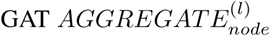 function in more details in the Results section.

Regarding the *COMBINE*_*graph*_ function, combining graph-level embeddings at different levels is relatively not a concern since GNNs rarely consider more than two layers. Kipf et al. [37] reported that considering more than two layers performs worse with their model. Although deeper models can in principle access to more information, some of the information may be “washed out” by too many aggregations.

Some studies alternatively considered either the sum [38, 43], the max or the concatenation [60] of the {**m**^(*l*)^} to define a final graph-level representation **m**. The concatenation stores explicitly sub-graph information at different granularity. Indeed, the aggregation process (at all nodes), after *l* iterations, learns the topology as well as the distribution of node features in the “*l* + (*l* − 1)”-hop neighbourhood. It also allows the network to skip the last layers if they happen to be harmful for the prediction. Xu et al. [60] also proposed a biLSTM-attention module processing the sequence (**m**^(1)^, …, **m**^(*L*)^), but it may overfit on small graphs due to its relatively high complexity.

### 3.3 Protein sequence encoder

Before encoding a protein, i.e. a sequence of amino acids, with neuron network-based encoders, we need to define attributes for amino acids.

An obvious solution is to use the so-called “one-hot” encoding, where amino acids of the protein sequence are encoded by a binary vector of the length 20 (one bit for each natural amino acid), containing a single one and 19 zeros.

However, richer encodings are available. For instance, an amino acid can be encoded by a mutation probability profile. This position-dependent probability profile corresponds to the probability of the current amino acid to be mutated in other amino acids. It is usually calculated by running sequence alignment algorithm like PSI-BLAST [61] against a reference database. Additional structural or physical amino acid properties can be used to complete amino acid attribute vectors. However, this knowledge is not available at the proteome level, and therefore, in the context of chemogenomics, one cannot rely on such properties.

Because a protein is a sequence of amino acids, their appear very well suited to representation learning methods based on convolutional neuron networks (CNN) and recurrent neuron networks (RNN), in particular Long Short-Term Memory cells (LSTM) and bi-directional LSTM (bi-LSTM). Indeed, these neuron network architectures are dedicated to encode data of sequential nature. Indeed, they have been successfully designed and trained on proteomic data to achieve state-of-the-art performance in various prediction problems [62, 63, 64, 65, 66, 67].

Intuitively, one-dimensional convolutional layers can efficiently process protein sequences by detecting local patterns. Indeed, CNN filters provide a learnable bank of sequence motifs. Bi-directional LSTM (or RNN in general) layers can integrate local information both forward and backwards in the sequence, to encode local and contextual information. The local information could be the amino acid type if the bi-LSTM is built on the original description of amino acids, or amino acid patterns if the bi-LSTM is built on a convolutional layer.

### 3.4 Feed-forward neuron network

Feed-forward neuron networks (FNN), also called multi-layer perceptron (MLP), is the most common deep neuron network architecture. It consists of stacked fully connected layers, such that each neuron at a layer *l* takes as input all neuron outputs from layer *l-*1 and thus, directs its output to all neurons at layer *l* + 1 (which is what “fully connected” refers to). The original data are subjected to non-linear transformations across several layers, such that intermediate layers can be viewed as intermediate hidden representations capture abstract representations of the original data. In this sense, such a network is a representation learning model, because the architecture is optimised so that the learnt representation is tuned to provide the best performance to the considered prediction task.

However, FNN can also directly use expert-driven representations as inputs, which has been reported to lead to state-of-the-art performances in many chemoinformatics applications, as detailed in the next sections. From a more general perspective, FNN take as input any description in the form of a numerical attribute vector, as in the chemogenomic neuron network presented above.

## 4 Related works

In the next two subsections, we briefly present some crucial published works in the field of chemoinformatics and chemogenomics with deep learning methods.

### 4.1 Deep learning for chemoinformatics

Deep learning in chemoinformatics refers to prediction problems involving molecules (such as predicting solubility or toxicity), but not to the prediction of protein-ligand interactions as in chemogenomics. Therefore, chemoinformatics deep learning only involves two of the four blocks presented in Fig. 1: the molecule encoder and the prediction block which consists in an *MLP*_*pair*_ (i.e. FNN) in most studies.

The striking arrival of deep neuron networks into the field of DTI prediction occurred during the virtual screening Merk challenge whose winners used FNN on top of expert-based features. In later studies [68], the winners of the challenge performed an exhaustive evaluation of the effect of hyper-parameters of FNNs to predict binding of around 140,000 compounds in 19 target assays from the PubChem database. They encoded molecules with 3000 expert-driven structural and physicochemical descriptors that are calculated by the DRAGON toolkit [69]. Note that in this challenge, the prediction methods were single-task “ligand-based” prediction methods, one task corresponding to the prediction of ligands for a given protein. This ligand-based approach would not have been applicable to tackle the question of ligands specificity, which would have required to build a prediction model for each protein, at the proteome level, as explained in the Introduction section.

A few studies have explored multitask prediction on various chemoinformatics prediction tasks with deep neural networks. Their principle is that the prediction tasks share the same input and hidden layers, but each task has its own output node. In that context, Ramsundar et al. [70] used several datasets containing information about toxicity, pitfalls in virtual screening, results of biological assays, or interactions with protein targets. They reported that multitask FNN outperforms other approaches, and that adding more training data was still improving the multitask model. However, Dahl et al. [68] showed that multitask FNN performed slightly better than singletask FNN only for some tasks. This was confirmed by Xu el al. [71] who showed that positive transfer learning is conditioned by sharing a significant amount of input data whose labels are correlated. In general, other studies [72, 73, 74, 75] reported that FNN performs the best for a variety of chemoinformatics problem, but only for some metrics and datasets.

Overall, it is however possible to consider that FNN, with expert-driven chemical descriptors as inputs, is a reference method in deep learning for chemoinformatics problems, to which other methods should be compared.

Instead of using expert-driven chemical descriptors as input of the FNN, other studies used Recurrent Neuron Networks (RNN) modules processing SMILES representations of molecules, or Graph Convolution Networks (GNN) [38, 46, 75] processing the molecular graph, to learn descriptors for the molecules. RNN and GNN learn molecule descriptors respectively from the SMILES representation and the molecular graph in such a way that the feature are optimised to maximise the objective function. Approaches based on RNN modules processing SMILES proved to be competitive with GNN which directly process the molecular graph [32, 33].

However, GNNs is a very active field in deep learning. They display valuable extension repertoires, and their flexibility offer a larger room for improvements, as presented above. Therefore, we considered GNN in the present work, as detailed in the following.

### 4.2 Deep learning for chemogenomics

In chemogenomics, where interaction predictions are made for many molecules and proteins within the same model, one need to encode proteins and ligands, combine these representations, and use them as input of the neuron network that calculates the outputs, as represented in Fig. 1. As in chemoinformatics, the protein and molecular descriptors can either be calculated based on expert-based toolkits, or learnt with an encoder.

In the case of learnt representations, molecules descriptors are learnt from molecular SMILES or graph representations, respectively with RNNs or GNN, as in chemoinformatic approaches presented above. Protein descriptors are learnt from the protein sequence of amino acids with a recurrent neuron network (RNN) encoder, or with a 1D Convolution Neuron Network (CNN) encoder. The molecule and protein representations are combined to build a pairwise latent representation used as input of a FNN, in order to predict whether the molecule and the protein interact.

Direct predecessors of the chemogenomic neuron network that we investigate in the present study are introduced below.

Ozturk et al. [76] used the following architecture for the four blocks of the chemogenomic neural network: (1) a three layers CNN on SMILES repreŝentations to encode molecules, (2) three layers CNN on protein sequences to encode proteins, (3) concatenation of the learnt molecule and protein encodings to combine them, which is a first natural attempt to combine the molecule and protein descriptors, and (4) a three layers FFN to predict the affinity value between the tested molecule and protein.

The authors considered the Davis and Kiba datasets, both containing selectivity assays in the kinase protein family, so that this study focuses on DTI prediction within a single protein family. The Davis dataset contains interactions between 442 proteins and 68 ligands and the Kiba dataset contains interactions between 467 targets and 52 498 drugs. They obtained better or similar performance compared to Simboost [77] (a gradient boosting machine-based method) and KronRLS [16] (analogue of KronSVM but with kernelised RLS) in terms of Mean Square Error (MSE). When “binarying” the outputs (they set the threshold of pKd value, distinguishing actives from inactives, to 7), their method performed the best, in terms of AUPR, for both datasets, but significantly better only for the KIBA dataset. Importantly, they did not compare to FNN (with expert-driven descriptors for molecules and proteins as inputs) and to *NRLMF*, which led to state-of-the-art performances in many studies, as mentioned previously.

In a more sophisticated approach, Tsubaki et al. [78] considered the following blocks: (1) GNN modules to encode molecules, in which the molecular graph-level representation **m**^(*l*)^ is learnt with an aggregation function 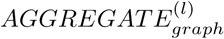 corresponding to the sum over the atom level representations, as in the minimal GNN defined above. (2) Protein encoding **h**_*prot*_ is performed with a CNN on amino acid representations which learn latent representation at the amino acid-level 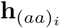. (3) The *Comb* module uses an attention mechanism that affects weights to learnt representation of each amino acid in the sequence. These weights are learnt via the similarity between a learnable linear transformation (parameterised by **W**_*inter*_ and **b**_*inter*_) of the molecular graph-level and amino acid-level representations:

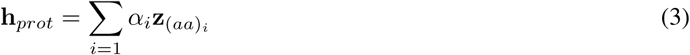

in which 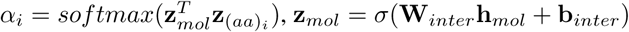, **z**_*mol*_ = *σ*(**W**_*inter*_ **h**_*mol*_ + **b**_*inter*_) and 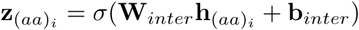.

In the previous equations, *σ* stands for the sigmoid function and softmax is the standard softmax function defined on n-dimensional vectors by: 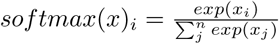

Let us put the above equations into words: the more the representation of a given amino acid *aa*_*i*_ is similar to that of the molecule (after the linear transformation), the higher its weight *α*_*i*_ in the weighted sum used to define the protein representation **h**_*prot*_. An intuitive interpretation of the weight *α*_*i*_ is that they indicate what parts of the protein sequence mostly govern individual predictions.

This attention mechanism allows interpretation of predictions based on which amino acids are involved in ligand binding. This was tested by comparing the amino acids found to be critical for interaction with a molecule by the attention mechanism to those found to interact with the molecule when docking the ligand into the binding pocket of the protein. Note that this control is based on a prediction methods (docking), and not on experimental data, and that it can be performed only for proteins with known 3D structures.

The authors used two datasets extracted from the DrugBank database. First, a human data set with 3,500 positive interactions between 1,000 unique molecules and 850 unique human proteins. Second, a C.Elegans dataset containing 4,000 positive interactions between 1,434 unique molecules and 2,504 unique C.Elegans proteins. They compared the performance of their method to an approach based on SVM with fingerprint-RBF kernel (not known to be state-of-the-art comparing to the use of the Tanimoto kernel), on various test “positive:negative” ratios (1:1, 1:3 and 1:5). Indeed, much more negative than positive interactions are recorded, and considering more negative interactions in the test set correspond to a more realistic setting in the context of proteome-wide DTI prediction.

For the human-specific dataset, the precision score is always higher for SVM (around 5% improvement), but conversely, the ROCAUC is always higher for their deep learning approach (from 1% to 5% improvement). Overall, the ROCAUC varied in the range of 0.91 to 0.97. For the recall score, SVM suffers more from an imbalanced positive:negative ratio than their deep learning approach.

In conclusion, the deep learning method or the SVM-based method performs better depending on the metrics that is considered.

Although this study paved the way for deep learning in chemogenomics, going one step further than Ozturk et al. [76] in terms of deep learning technology, it suffers from several lacks in the discussion of their approach. First, the authors did not compare the performance of their method to those obtained without the attention mechanism, i.e. simply encoding protein sequences with a CNN, which limits interpretation of their approach. Importantly, they did not compare their method to the simpler reference FNN with expert-based descriptors as inputs, which we will see below in the Results section, often leads to the best prediction performances.

In addition, it does not provide a clear comparison with state-of-the-art DTI prediction without deep learning. Indeed, their results on the gold standard Yamanishi dataset [13] display similar performance than kronRLS and KBMF2K [29] (a probabilistic matrix factorisation approach), while these methods were shown to be outperformed by *NRLMF* on this dataset. Moreover, the choice of the Yamanishi dataset is questionable here. Indeed, it is relatively small, so that it is not well suited to fairly evaluate deep learning models, as will be further discussed in the Results section.

Overall, we are far from the striking gap of performance obtained by deep learning-based models in the image processing and natural language processing fields. Moreover, the datasets considered in the above studies are very heterogeneous, and recent promising developments in GNN technologies have not yet been explored in chemoinformatics, and in particular, in chemogenomics.

Therefore, in the present paper (see Results section), one of the directions that we explored is related to some of the most promising directions in GNN research, to evaluate their potential to improve predictions with the chemogenomic neuron network.

In the following, to investigate whether neuron networks and related recent developments can lead to significant improvements for chemogenomics, we first compare, on the two DrugBank-based datasets, the standard formulation of the chemogenomic neuron network with the state-of-the-art methods presented above.

Indeed, the chemogenomic neuron network is very modular, which let significant room of improvement. Therefore, we then evaluate separately promising neuron network architectures for the molecular graph encoder, protein sequence encoder and *Comb* operation, in order to explore various deep learning strategies for data featurisation.

Then, we adopt another perspective to improve chemogenomic predictions with deep learning. Indeed, data featurisation and data augmentation techniques are independent levers of improvement for the chemogenomic neuron network prediction performance. Therefore, we also explore the simultaneous use of expert-based and learnt features in a differentiable fashion, with the idea that combining both types of features might increase performances. This strategy uses multiple views of the data as a way to provide “more informative” data. We also explore data augmentation thanks to transfer learning via curriculum learning. This refers to using an additional larger dataset for pre-training the neuron network with an auxiliary task, before using them it in the chemogenomic task of interest, with the aim of learning richer representations of molecules or proteins. We tested various way to implement this idea.

## 5 Materials and Methods

### 5.1 Dataset

Several bio-activity databases are publicly available. Some provide quantitative measurements such as the IC50, EC50, Ki and potency (in particular the PubChem [79], ChEMBL [80] and BindingDB [81] databases), while others report binary information on activities or interactions between molecules and protein pairs, like the DrugBank [82] or KEGG [83] databases.

The DrugBank database [82] is a widely considered bio-activity database. Although much smaller than PubChem and ChEMBL, it provides high-quality information for approved and experimental drugs along with their targets. It contains around 17,000 curated drug-target associations with high-quality standards.

Therefore, we used the DrugBank database version 5.1.0 to build two druggable proteome-wide drug-target interaction datasets. In the first dataset, called *DBH*, we only kept interactions including human proteins and their ligands, whereas in the second, called *DBEC*, we only kept interactions including Escherichia Coli proteins and their ligands.

*DBEC* is composed of 592 molecules targeting a total of 314 proteins, and includes 874 protein-ligand interactions. *DBH* is composed of 4834 molecules targeting a total of 2561 proteins, and includes 13070 protein-ligand interactions. In both cases, recorded protein-ligand interactions constitute the positive training pairs, and the vast majority of the targets and drugs are involved in only one or two known interactions.

All other protein-ligand pairs are unlabeled because no interactions were recorded for them in the database. Most of these pairs are expected not to interact, but a small number of them are in fact missing interactions. However, as an approximation, we considered that all unlabeled pairs as negative examples.

As we later examine curriculum learning to increase the performance on these DrugBank-based datasets, we also consider a PubChem-based and a protein sub-cellular localisation datasets as auxilliary datasets for neuron network pre-training.

We pre-processed the PubChem datasets included in the MolNet dataset collection [84] (namely, the PCBA dataset) to keep the bioassays composed of a maximum of common molecules. The resulting dataset consists in a total of 439, 863 unique molecules and 90 binary classification tasks (bioassays). For each bioassay, molecules are labelled as active, inactive or unknown. When a molecule is labelled as unknown for a bioassay (because it has not been experimentally tested), the prediction error for the corresponding task and molecule is set to zero.

The sub-cellular localisation dataset [67] is a multi-class classification problem, labelling 5917 human proteins with one of the twelve sub-cellular localisation considered as classes.

### 5.2 Evaluation procedures

We evaluated the performance by 5-fold nested cross-validation. Although we recorded the ROCAUC [85] and AUPR [86] scores for each test fold, we predominantly refer to the AUPR. Indeed, the AUPR score is considered as a more significant quality measure than the ROCAUC when negative interactions are in fact unknown interactions (which is the case in DTI prediction), and when there are many more negative than positive samples in the test set (which is also the case in DTI prediction).

Indeed, the AUPR score emphasises on the recovery of the positive samples and penalises the presence of false positive examples among the best-ranked samples. Therefore, we used AUPR as the evaluation metric for the hyper-parameters optimisation and in the early stopping procedure used in deep learning approaches.

First, we considered four way to split our Drug-Bank-based dataset for the cross-validation scheme, each corresponding to a typical chemogenomic setting. The *S*_1_ settings corresponds to data split at random, *S*_2_ to the orphan ligand case (pairs in one fold only contain molecules that are absent from all other folds), *S*_3_ to the orphan protein case (pairs in one fold only contain proteins that are absent from all other folds), and *S*_4_ to double orphan (pairs in one fold only contain proteins and molecules that are both absent from all other folds). This approach was suggested by Pahikkala et al. paper [87], because it allows assessment of performance in various real-life situations in chemogenomics. Note that the folds of *S*_4_ were built by intersecting those of *S*_2_ and *S*_3_, so that there are 25 folds in the *S*_4_ setting instead of 5.

Independently, we considered scenarios in which the “positive:negative” sample ratio in the test set is 1:1, 1:2 or 1:5., in order to evaluate whether methods remain robust when predicting interactions at the proteome level (i.e. when there are much more non-interacting than interacting pairs).

### 5.3 Reference similarity-based approaches

As reference methods not involving deep learning, we considered the kronSVM [7] and *NRLMF* [11] approaches as they led to state-of-the-art performances, in particular on the gold standard Yamanishi dataset. These methods have been shortly introduced in section 2 of the present study.

For kronSVM, the regularisation hyper-parameter (often called C) of SVM was automatically optimised when assessing the performance via 5-fold nested cross-validation. We set the five hyper-parameters of *NRLMF* to their default value, as stated in the original paper [11].

These two approaches use molecule and protein valid kernels.

For proteins, we used the local alignment kernel [88] (LAkernel) that mimics the behaviour of the Smith-Waterman (SW) score. While the SW score only keeps the contribution of the best local alignment between two sequences to quantify their similarity, the LAkernel sums up the contributions of all possible local alignments. LAkernel has been shown to overtake other protein sequence kernels on protein homology detection [88]. This kernel compares proteins from the evolutionary point of view, but it seem relevant for drug virtual screening because a local similarity of sequence may correspond to a local similarity of binding pockets [28].

For molecules, we used the Tanimoto similarity measure, defined as the ratio between the number of substructures shared by the two molecular graphs over the number of all substructures considered. This similarity measure is very widely used in chemoinformatics, and it is also a valid kernel [89]. Moreover, ECFP4 fingerprints of molecules have been reported to perform the best [90] and are widely used for drug virtual screening.

Therefore, we also considered the Tanimoto metric on the Morgan fingerprints (analogue of ECFP4) as the molecule kernel, that we computed thanks to RDKit library [91] in Python.

### 5.4 Reference feed-forward neuron networks

As motivated in section 4, we considered as reference method in deep learning, the feed-forward neuron networks (FNN). This reference method consists in an FNN that uses the concatenation of expert-based numerical feature vectors as input, for proteins and molecules in our case.

Therefore, we extracted 1021-dimensional structural Morgan fingerprint vectors (analogues of ECFP4) with the RDKit library [91] for each molecule. Indeed, when used individually, ECFP4 fingerprints have been reported to perform the best [90] and are widely used for virtual screening.

For protein descriptors, with the ProtR package, we extracted 1920-dimensional feature vectors corresponding to 11 protein level feature groups including amino acid composition (AAC), dipeptide composition (DC), autocorrelation descriptors, “composition, transition and distribution” descriptors (CTD), quasi-sequence-order descriptors (QSO), pseudo-amino acid composition descriptors (Pse-AAC), amphiphilic pseudo-amino acid composition descriptors (Am-Pse-AAC), topological descriptors at atomic level and total amino acid properties. In general, these descriptors encode the fraction of each amino acid type and amino acid pair type within a protein as well as the distribution of amino acid properties along the sequence. They also encode the composition, transition and distribution of attributes (including hydrophobicity, normalised van der Waals volume, polarity, and polarisability) along the sequence. Indeed, Ong et al. [92] reported that combining these descriptors provide the best results, for the prediction of functional families with support vector machines, although none of them performed significantly better than the others individually.

We “manually” optimised the hyper-parameters of FNN (number of layers, number of neurons in each layer, initial learning rate, batch size, learning rate decay factor, dropout probability and weight decay) on a fixed train/validation/test data split. Notably, we spend little time in fine-tuning these hyper-parameters, compared to those of the chemogenomic neuron network.

We set the number of neurons to 2000, 1000 and 100 for the successive three stacked fully connected layers, weight decay and dropout probability to 0 (no regularisation), the initial learning rate to 10^*-*3^, the batch size to 100, the learning rate decay to 0.9, the number of epochs (number of times the algorithm sees the entire dataset) to 100 and early stopping patience to 20. The early stopping patience is the number of successive epochs, for which the performance on the validation data decreased, required to stop training.

### 5.5 The “standard” chemogenomic neuron network

The main approach we investigate in this study is the chemogenomic neuron network that we introduced in the previous sections.

This study is meant to evaluate the potential of promising modifications of the neuron network architecture and appealing data manipulation, compared to what we define as our “standard” chemogenomic neuron network formulation.

In this standard formulation, the encoder for molecules is the “minimal” GNN, defined in section3 by the Alg. 1, in which the 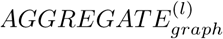 consists in the sum function, 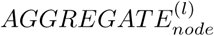 consists in eq. 2, and the *COMBINE*^*graph*^ function is set such that the last graph-level embedding **m**^(*L*)^ is used as the graph-level representation **m**.

The GNN requires original atom and bond features as input. We used the RDkit library to extract the following features (based on Coley et al. work [43]): “atom identity”, “number of hydrogens”, “number of heavy neighbours”, “formal charge”, “is in ring”, “is aromatic”, “polar surface area”, “partial charge”, “chirality tag”. The chemical bond description contains: “bond type”, “is in ring”, “is aromatic”, “is conjugated”.

In our “standard” chemogenomic neuron network, the protein sequence encoder is a stacked 1D convolutional neuron layers (CNN), with amino acids described by their types in a one-hot encoding fashion. The final representation learnt for the protein is the sum of the learnt amino acid representations resulting from the stacked convolutional layers.

Finally, the *Comb* operation, in the “standard” formulation, is the concatenation of the molecule and protein learnt representations.

Compared to the architecture proposed by Tsubaki et al. [78], the standard chemogenomic neuron network described above does not use any bonds attributes for the molecule, and does not use any attention mechanism.

We also tested more complex architectures derived from the standard chemogenomics network. These architectures are described directly in the Results section.

We optimised three sets of hyper-parameters (architecture, learning and regularisation hyper-parameters) by recording the performance on multiple train/validation/test data splits to explore hyper-parameter spaces manually, since a grid-search or random-search would have required too much calculation time. In the following, we detail the optimised hyper-parameters and the range of values we considered are given in parentheses.

The architecture hyper-parameters are the number of filters both for the molecule and protein encoders ([10, 1000]), the number of layers for both the protein and molecule encoders (⟦2, 5⟧), the convolutional filter size ([6, 12]) and the convolutional stride for the sequence encoder ([2, 6]), and the number of neurons in the last hidden fully connected layer which processes the pairwise-level encoding ([10, 1000]).

The learning hyper-parameters are the batch size ([1, 100]), initial learning rate ([10^*-*5^, 10^*-*2^]) and learning rate decay factor ([0.8, 0.99]).

The regularisation hyper-parameters are the probability of dropout ([0., 0.9]) and the weight decay ([10^*-*4^, 0.1]). We set the number of epochs to 100, and we also considered early stopping such that training stops if the performance (AUPR score) on the validation set does not increase in ten successive epochs. In practice, training was always stopped early.

After exploring the hyper-parameter spaces, we set, for both datasets, the number of filters to 100, the number of convolutional layer to 3, the stride to 3, the filter size to 8, the weight decay and dropout to 0, batch size to 20, initial learning rate to 10^*-*3^, learning rate decay factor to 0.9 and number of neurons in the last prediction layer to 100. We found that training was not sensitive to relatively small variations of hyper-parameters. In particular, adding a small quantity of dropout (until 0.5) leads progressively to a longer training and a loss of training performance, whereas it did not significantly improve the performance on the validation data.

Furthermore, we repeated training and testing procedures of the chemogenomic neuron network three times for *DBEC* and two times for *DBH* (because the training was computationally too expensive), and kept only the best achieved performance. Indeed, we noticed that the performance of the chemogenomic neuron network varied significantly depending on the initial random seed.

## 6 Results and discussion

### 6.1 Comparison of the “standard” chemogenomic neuron network to reference methods

We first compared the performance of the standard chemogenomic neuron network (CN) to those of the considered state-of-the-art reference methods, namely SVM and *NRLMF* for machine-learning methods not involving deep learning, and FNN for deep-learning methods, as described in the Materials and Methods section. Performances are reported in Tables 1, 2, 3 and 4, displayed in the format “mean score ±score standard deviation”, for various ratios of positive:negative test samples.

**Table 1:** ROCAUC scores on the *DBEC* dataset in the four settings and for a test sample positive:negative ratios in {1:1, 1:2, 1:5}

| | raw ($S_1$) | | | orphan proteins ($S_2$) | | | orphan molecules ($S_3$) | | | double orphan ($S_4$) | | |
| --- | --- | --- | --- | --- | --- | --- | --- | --- | --- | --- | --- | --- |
|  | 1:1 | 1:2 | 1:5 | 1:1 | 1:2 | 1:5 | 1:1 | 1:2 | 1:5 | 1:1 | 1:2 | 1:5 |
| SVM | **75.14 ± 1.7** | 74.95 ± 2.04 | 74.99 ± 1.76 | 54.97 ± 4.33 | 56.5 ± 4.29 | 56.67 ± 4.44 | 74.03 ± 3.8 | 73.89 ± 3.78 | 74.14 ± 3.72 | 53.84 ± 8.86 | 54.46 ± 8.43 | 53.06 ± 9.63 |
| *NRLMF* | **78.8 ± 3.68** | **78.88 ± 2.73** | **78.41 ± 2.69** | **65.05 ± 7.1** | **64.96 ± 6.32** | **65.4 ± 4.9** | **77.19 ± 1.65** | **77.21 ± 2.06** | **77.38 ± 1.81** | **65.2 ± 7.26** | **64.59 ± 6.26** | **63.61 ± 6.51** |
| FNN | **75.92 ± 1.17** | 74.63 ± 1.04 | 74.88 ± 1.52 | **58.25 ± 5.34** | 55.12 ± 4.67 | 54.89 ± 4.98 | 74.67 ± 2.38 | **75.52 ± 2.4** | 73.96 ± 3.15 | 56.03 ± 9.22 | 55.91 ± 8.07 | 54.68 ± 8.12 |
| chemogenomic neuron network | **77.1 ± 1.83** | **76.5 ± 0.89** | **76.74 ± 0.85** | **61.9 ± 4.41** | **59.64 ± 8.55** | **64.25 ± 4.0** | 73.49 ± 3.99 | 74.04 ± 4.91 | 74.91 ± 3.98 | **59.23 ± 9.5** | **58.4 ± 8.74** | **58.62 ± 9.1** |

**Table 2:** AUPR scores on the *DBEC* dataset in the four settings and for a test sample positive:negative ratios in {1:1, 1:2, 1:5}

| | raw ($S_1$) | | | orphan proteins ($S_2$) | | | orphan molecules ($S_3$) | | | double orphan ($S_4$) | | |
| --- | --- | --- | --- | --- | --- | --- | --- | --- | --- | --- | --- | --- |
|  | 1:1 | 1:2 | 1:5 | 1:1 | 1:2 | 1:5 | 1:1 | 1:2 | 1:5 | 1:1 | 1:2 | 1:5 |
| SVM | **81.75 ± 1.75** | **73.25 ± 2.4** | **61.55 ± 3.11** | 59.62 ± 5.55 | **45.89 ± 6.73** | **28.95 ± 7.66** | **80.79 ± 2.37** | **72.02 ± 3.06** | **60.96 ± 3.73** | 58.52 ± 9.21 | 43.37 ± 9.15 | 24.4 ± 8.8 |
| *NRLMF* | **82.56 ± 2.82** | **73.89 ± 2.98** | **59.89 ± 4.35** | **67.56 ± 7.04** | **53.18 ± 8.12** | **35.62 ± 8.07** | **81.49 ± 1.48** | **72.35 ± 2.2** | **60.06 ± 2.73** | **67.93 ± 8.63** | **52.97 ± 8.7** | **34.5 ± 9.75** |
| FNN | **79.82 ± 0.98** | 68.07 ± 2.35 | 51.55 ± 2.55 | **61.92 ± 5.11** | 42.24 ± 5.27 | 24.26 ± 3.91 | 78.57 ± 2.22 | 68.14 ± 3.43 | 49.34 ± 3.11 | **60.04 ± 8.83** | 43.55 ± 9.84 | 22.97 ± 6.59 |
| chemogenomic neuron network | 79.04 ± 2.24 | 66.5 ± 2.51 | 49.08 ± 4.31 | **60.79 ± 6.24** | **45.11 ± 8.65** | **28.38 ± 5.81** | 76.23 ± 2.23 | 64.61 ± 4.78 | 46.14 ± 4.92 | **61.97 ± 8.72** | **45.81 ± 8.5** | **27.0 ± 6.96** |

**Table 3:** ROCAUC scores on the *DBH* dataset in the four settings and for a test sample positive:negative ratios in {1:1, 1:2, 1:5}

| | raw ($S_1$) | | | orphan proteins ($S_2$) | | | orphan molecules ($S_3$) | | | double orphan ($S_4$) | | |
| --- | --- | --- | --- | --- | --- | --- | --- | --- | --- | --- | --- | --- |
|  | 1:1 | 1:2 | 1:5 | 1:1 | 1:2 | 1:5 | 1:1 | 1:2 | 1:5 | 1:1 | 1:2 | 1:5 |
| SVM | 74.6 ± 0.57 | 74.6 ± 0.55 | 74.62 ± 0.54 | 50.26 ± 3.05 | 49.88 ± 3.06 | 49.88 ± 2.95 | 75.16 ± 1.51 | 75.28 ± 1.76 | 75.14 ± 1.69 | 49.79 ± 1.61 | 50.25 ± 1.84 | 50.5 ± 1.68 |
| *NRLMF* | 82.98 ± 0.55 | 83.06 ± 0.65 | 83.12 ± 0.63 | 65.18 ± 1.36 | 65.18 ± 1.28 | 65.23 ± 1.2 | 59.49 ± 0.96 | 59.56 ± 0.81 | 59.6 ± 0.93 | 50.37 ± 1.48 | 50.29 ± 1.55 | 50.47 ± 1.35 |
| FNN | **90.39 ± 0.42** | **90.65 ± 0.46** | **90.45 ± 0.52** | 74.74 ± 0.4 | 74.08 ± 2.29 | 74.25 ± 1.96 | 83.06 ± 1.48 | **83.52 ± 1.52** | 82.9 ± 1.8 | **69.12 ± 2.3** | **69.17 ± 1.99** | 68.79 ± 1.69 |
| chemogenomic neuron network | 88.67 ± 1.59 | 88.21 ± 1.47 | 88.21 ± 2.23 | **81.5 ± 1.47** | **80.83 ± 2.7** | **80.28 ± 1.4** | **85.76 ± 1.5** | **85.4 ± 2.02** | **85.76 ± 2.23** | **73.34 ± 5.89** | **71.21 ± 5.53** | **73.68 ± 4.87** |

**Table 4:** AUPR scores on the *DBH* dataset in the four settings and for a test sample positive:negative ratios in {1:1, 1:2, 1:5}

| | raw ($S_1$) | | | orphan proteins ($S_2$) | | | orphan molecules ($S_3$) | | | double orphan ($S_4$) | | |
| --- | --- | --- | --- | --- | --- | --- | --- | --- | --- | --- | --- | --- |
|  | 1:1 | 1:2 | 1:5 | 1:1 | 1:2 | 1:5 | 1:1 | 1:2 | 1:5 | 1:1 | 1:2 | 1:5 |
| SVM | 78.34 ± 0.44 | 66.85 ± 0.68 | 48.99 ± 0.5 | 50.35 ± 2.64 | 33.46 ± 2.33 | 16.82 ± 1.54 | 78.2 ± 1.77 | 66.86 ± 2.12 | 48.66 ± 2.74 | 49.84 ± 1.7 | 33.26 ± 1.64 | 16.81 ± 1.02 |
| *NRLMF* | 86.14 ± 0.45 | 78.73 ± 0.75 | 66.93 ± 1.06 | 68.19 ± 1.55 | 53.68 ± 1.72 | 34.75 ± 1.65 | 61.88 ± 0.99 | 45.93 ± 0.72 | 26.42 ± 0.52 | 50.45 ± 1.31 | 33.71 ± 1.25 | 16.99 ± 0.8 |
| FNN | **91.45 ± 0.45** | **86.5 ± 0.44** | **75.91 ± 0.49** | 78.16 ± 0.44 | 66.02 ± 1.86 | **49.13 ± 2.58** | **85.69 ± 1.05** | **78.15 ± 1.18** | **64.06 ± 1.6** | **71.46 ± 2.02** | **57.93 ± 2.7** | **38.61 ± 2.29** |
| chemogenomic neuron network | 88.74 ± 1.68 | 80.29 ± 2.65 | 65.46 ± 4.53 | **81.82 ± 1.9** | **70.39 ± 3.66** | **47.68 ± 4.38** | **86.0 ± 1.85** | **77.02 ± 3.41** | 62.22 ± 4.66 | **72.26 ± 6.42** | **55.98 ± 7.67** | **38.48 ± 7.5** |

Figures 2 and 3 display the ROCAUC and AUPR performance obtained on the *DBEC* and *DBH* datasets in the case of a “positive:negative” test sample ratio set to 1: 5.

**Figure 2:**
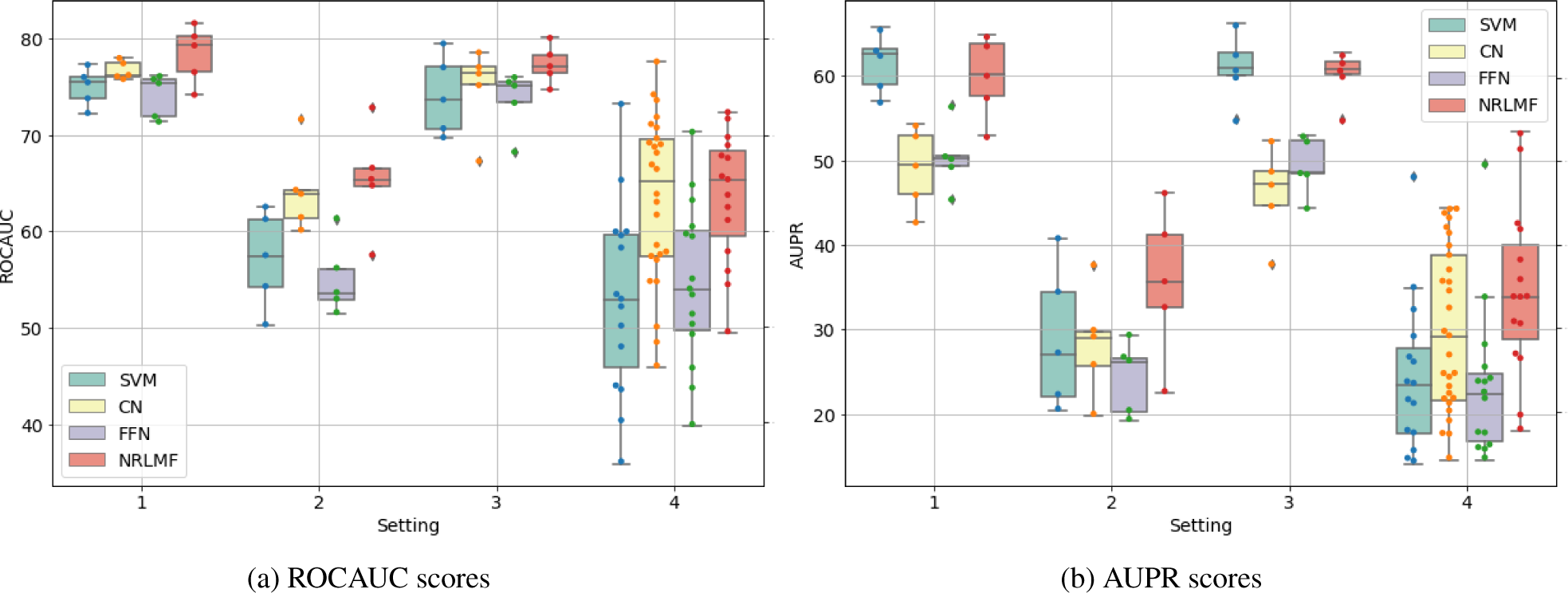
ROCAUC and AUPR performance on *DBEC* for the four *S*1, *S*2, *S*3, and *S*4 settings with a test and train sample positive:negative ratio set to 1:5. For each setting, the order from left to right in which the results of the different methods are displayed corresponds to that of the legend in the corner of the figures.

**Figure 3:**
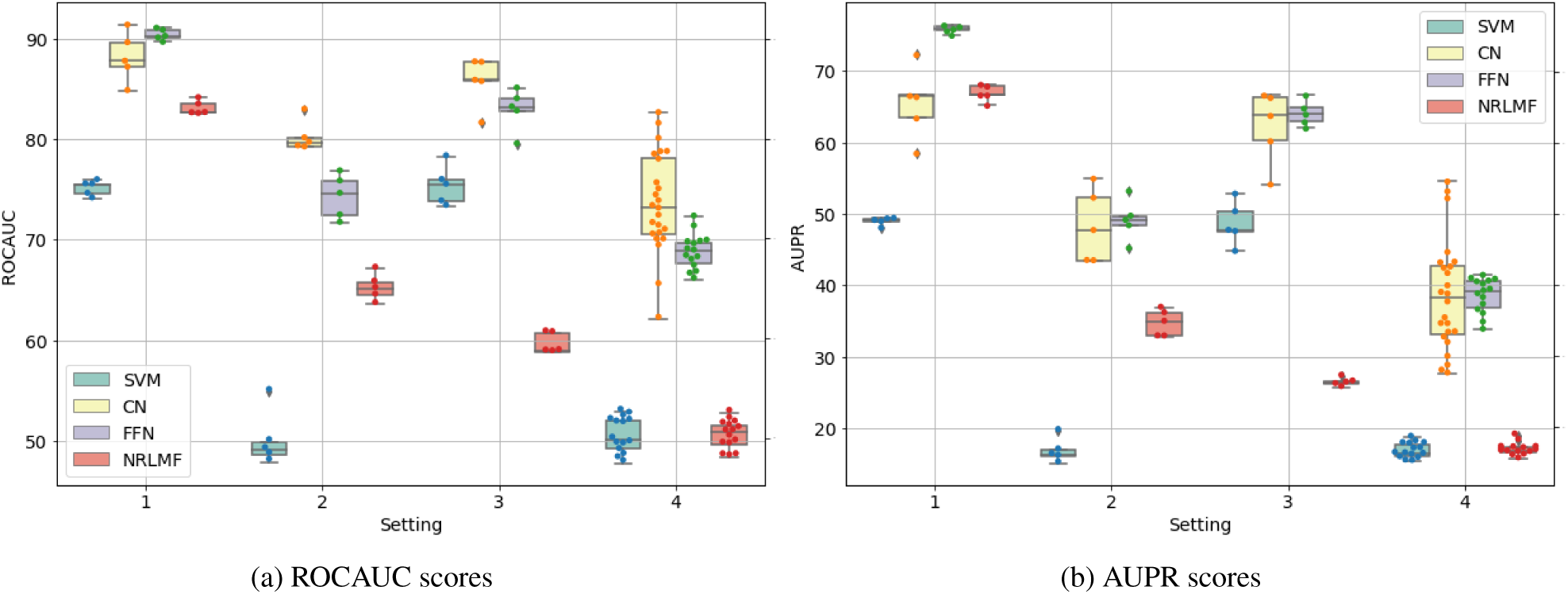
ROCAUC and AUPR performance on *DBH* for the four *S*1, *S*2, *S*3, and *S*4 settings with a test and train sample positive:negative ratio set to 1:5. For each setting, the order from left to right in which the results of the different methods are displayed corresponds to that of the legend in the corner of the figures.

To summarise the results, we can first notice that the “best” method depends on the dataset and the setting.

On the *DBEC* dataset, the *NRLMF* method performs globally better than SVM for non deep learning reference methods, and CN performs better than the reference method FNN for deep learning. Overall, the *NRLMF* method performs slightly better in all cases. This could be explained by the fact that *DBEC* is a relatively small dataset, which favours to the *NRLMF* and SVM approaches not involving deep learning.

On the larger *DBH* dataset, SVM outperformed *NRLMF* and CN outperformed FNN. Globally, the standard CN deep learning approach leads to the best results. It appears as a good default method that seems to be the most robust in a panel of settings.

Globally, the performance reached on the *DBH* are 10 points higher than those reached on *DBEC*, which was expected since there are much more data (more information) to train the models in *DBH* than in *DBEC*.

Furthermore, all methods appeared to be sensitive to an imbalanced test set. Indeed, when changing the “positive:negative” ratio in the test set from 1:1 to 1:5, all methods lost about 20% performance in AUPR on *S*_1_, and up to 30% on *S*_2_, *S*_3_, and *S*_4_, although SVM and *NRLMF* seem slightly more robust, as shown in Tables 1, 2, 3 and 4.

However, we expect that a relatively large number of drug-like molecules interact with more than one protein [93], whereas most of the molecules in the database have one or two known targets. Therefore, a significant amount of false predicted positives are expected to correspond to real interactions, which means that the performance in the case of the 1:5 “positive:negative” ratio, the performances might be underestimated.

Regarding the settings, the best scores are obtained for *S*_1_, the worst for *S*_4_, and the scores obtained in the *S*_2_ and *S*_3_ datasets are intermediate between those observed on *S*_1_ and *S*_4_. This was expected, since *S*_4_ corresponds to the double orphan test set, where no pair in the training set contains proteins or ligands present in the tested pairs to guide the predictions, whereas the models can rely on training pairs containing either the same proteins in *S*_2_ or the same ligands in *S*_3_.

The loss of performance between the raw (*S*_1_) and the double orphan (*S*_4_) settings is about 10 points of ROCAUC and 20 points of AUPR. More importantly, the prediction on *S*_4_ varies more depending on the test folds than in the other settings. This can be understood as a consequence of the small size of the test set in the *S*_4_ setting (any test fold is 1*/*25^*th*^ of the total amount of data whereas they represent 1*/*5^*th*^ in the other settings), which might result in more heterogeneous test sets.

Interestingly, the performance reached by all models is better for *S*_3_ than for *S*_2_, which suggests that predicting ligands for new protein targets is a more difficult task than predicting targets for new molecules. We think it is due to the bias in the data. Indeed, the DrugBank dataset from which our datasets were extracted, contains molecules from drug discovery projects that usually focused on some specific molecular scaffolds [94] binding to a given therapeutic target. Therefore, orphan molecules in *S*_3_ test sets will usually have similar molecules in the train set, because of this sampling bias in the chemical space, which is not true for orphan proteins in *S*_2_ test sets.

Let us conclude these analysis by a few remarks. We considered several “positive:negative” samples ratios in the train set, as all approaches may benefit from more negative training data. Overall, we tested four “positive:negative” train sample ratios (1:1, 1:2, 1:5 and 1:10). Recall that such “negative” training data are unknown interactions, so that we must not use of all of them to train the models. Moreover, it would be hardly computationally tractable.

We did not correct the imbalance proportions of labels in the training set for deep learning-based approaches, because we found that it damaged their final performance on the validation set.

The SVM and *NRLMF* methods did not benefit from an increasing number of negative samples in the train set, whereas deep learning-based methods, as expected, strongly did. More precisely, increasing the number of negative samples in the training set, up to five times the number of positive sample, increased the performance of CN and FNN by about 10% in AUPR (only 2% in ROCAUC) for the *DBH* dataset, and by 20% in AUPR (ten points of ROCAUC) for the smaller *DBEC* dataset. We also observed that CN benefits slightly less than FNN from an increasing number of negative in the training set. The performance of CN and FNN did increase between a ratio of 1:5 and 1:10 in some cases, meaning the neuron network-based models may still benefit more from more training negative samples.

In the following, we tested several changes in the “standard” chemogenomic neuron network (CN) discussed in the two preliminary studies described in the current section.

For all modifications considered in the following, we explored the associated hyper-parameter spaces to find the best reachable solution. To this end, we measured 5 times the performance on a single train/validation/test split for the *S*_1_, *S*_2_, *S*_3_ and *S*_4_ settings, since an evaluation of all of these modification by nested cross-validation was neither computationally feasible for us, nor relevant. The mean and standard deviation of the performance we obtained are reported in Table 5, and are discussed below. The performances are displayed in the format “mean score ±score standard deviation”.

**Table 5:**
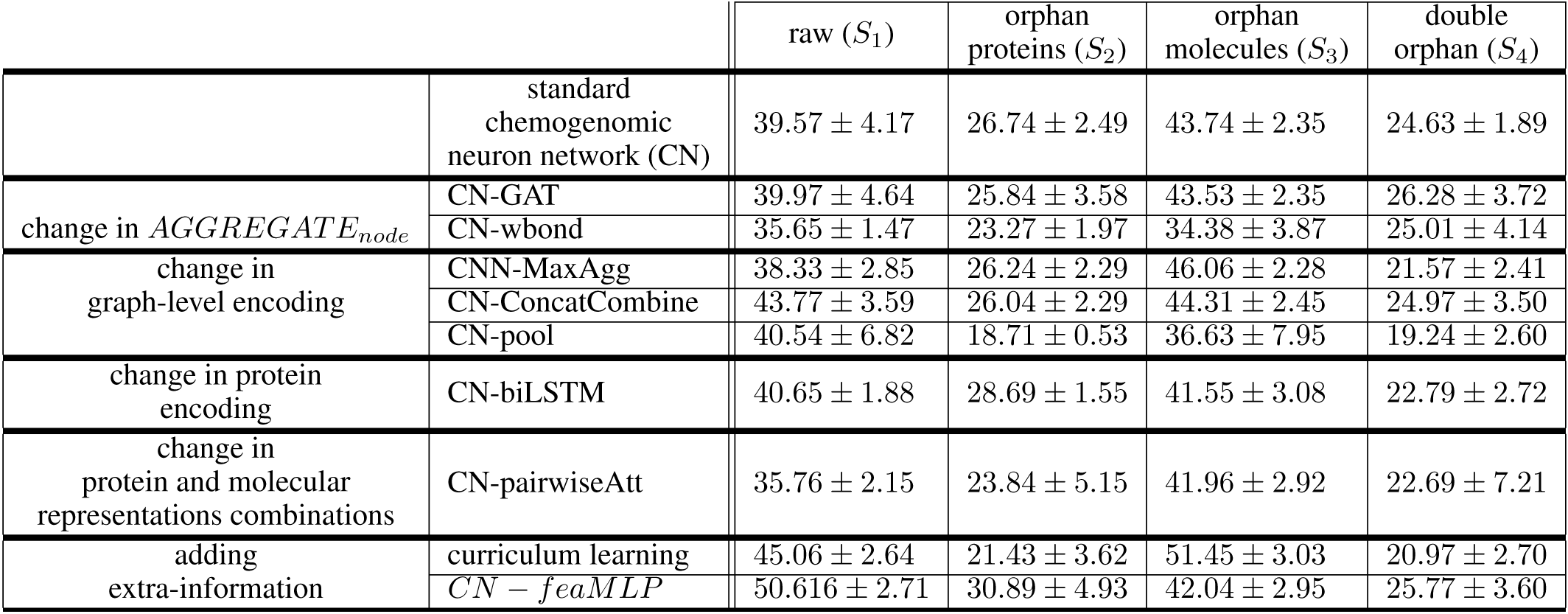
AUPR score of various modifications of the chemogenomic neuron network on a single train/validation/test split of the *DBEC* dataset. Standard deviations are obtained after repeating 5 times the evaluation procedure.

Building from the standard chemogenomic network studied in the present section, in the following sections, we evaluate modifications in the GNN architecture to encode molecules based on various nodes and graph aggregation functions and of various ways to combine molecule and protein representations, including two attention mechanisms.

### 6.2 Evaluation of molecular graph encoding architectures

In this section, we investigate promising modifications to the standard GNN formulation for molecules encoding. Among the modifications that have been proposed in the general field of graph representation learning, we only considered those that appeared the most relevant in the context of chemogenomics, both in terms of intuition and expected performance improvements, as discussed below.

Among 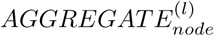 functions, the GAT function proposed by Velickovic et al. [58] seemed the most promising to us. Indeed, it resulted in significant performance improvement, and is intuitively relevant since it intends to prioritise relevant neighbours via a multi-head attention mechanism at each node update.

We also assayed the “wbond” function that takes into account the bond attributes for the aggregation. The aim was to test whether bond attributes can bring representation power and flexibility. Here, the “wbond” 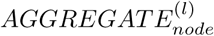 function refers to (as in Coley et al. [43]):

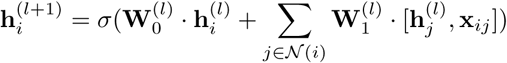

where *σ* is the sigmoid function, 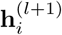 is the representation of atom *i* at step *l* + 1, 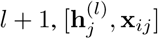 is the concatenation of the representation 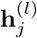 of atom *i* at step *l* and the bond attribute vector **x**_*ij*_ between atoms *i* and *j*, and 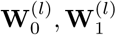,are learnable parameters.

We used the original implementation provided by the authors of the GAT method. For both GAT and “wbond”, we varied the number of convolutional filters (in {10,20,50,100,1000}) and the keeping probability of dropout (in {0,0.3,0.6}). We also tested several numbers of attention heads (in {1,5,10}) for the GAT function.

None of these two modifications lead to significant performance improvement for the four settings, as reported in Table 5.

Although the GAT function led to substantial improvements in gold standard datasets of graph representation learning, it did not in chemogenomics. This may be due to the the fact that molecular graphs are small, and that all atoms of molecules are informative when learning an end-to-end molecule encoding. In the case of the “wbond” function, the standard GNN may already leverage bond information based on the nodes attributes and degrees (i.e., based on the topology of the graph).

Independently, we tested several options to replace the summation in the 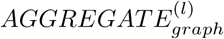 function and standard *COMBINE*_*graph*_ function of the minimal GNN, that builds the graph-level representation directly from node representations. From the simplest to the most sophisticated, we tested the max function [39, 60], the concatenation of graph level representation [60] and hierarchical differential pooling [51s, 52, 53], all introduced above.

When used as the 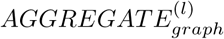 function, the sum function captures the distribution of node features in a single value. Alternatively, the max function is expected to capture representative elements in the node features’ distributions, with no additional parameters to learn. This could be relevant for molecular bio-activity prediction, as bio-activity may rely on representative elements such as pharmacophores or functional groups. The chemogenomic neuron network, whose GNN’s 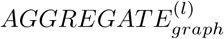 function is set to the max function, is mentioned as “CN-MaxAgg” in Table 5. Note that even if the sum function can theoretically capture representative elements, in particular when the dimension of the feature vector is high, the max function enforces capturing representative elements, which may improve the performance.

Regarding the *COMBINE*_*graph*_ function, we tested concatenation of graph-level representations after each updated **m**^(*l*)^, used to build the final graph-level representation **m**. We expect that it could help learning by skipping deleterious layers and by representing molecules via substructures of different sizes. The chemogenomic neuron network, whose GNN’s *COMBINE*_*graph*_ function is set to the concatenation function, is mentioned as “CN-ConcatCombine” in Table 5. Note that the simple sum function can, theoretically, adopt the same behaviour if the dimension of the feature vector is high enough, and if the network learns to store information in different locations in the feature vector at each update step.

The hierarchical pooling procedure [51, 52, 53] introduced earlier takes the role of both the 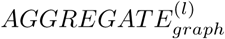 and *COMBINE*_*graph*_ functions. It seems particularly relevant as it is the analogue of the pooling operator in image processing. Indeed, it reduces the spatial information step by step by differentially coarsening the input graph after each node update step, until the graph is reduced to a single node. The chemogenomic neuron network whose GNN is based on hierarchical pooling is mentioned as “CN-pool” in Table 5. Again, note that a simple sum over node feature vectors can mimic the same behaviour, but the hierarchical pooling procedure is expected to enforce learning the graph-level representation in a hierarchical fashion and, hence, to improve encoding.

For all of these modifications of the molecular graph encoder, we varied the quantity of dropout ({0,0.3,0.6}). In the case of hierarchical pooling, that we implemented based on Ying et al. [51], we also varied the dimension of the nodes embedding (i.e. the number of convolutional filters) and the quantity of dropout when building the soft assignment matrices.

None of these modifications led to substantial performance improvements for the four settings on the *DBEC* dataset.

Intuitively, this means that, in the case of small chemical compounds, the simple sum function has the same representation power than the max for the 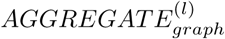 function and concatenation for the *COMBINE*^*graph*^ function. Therefore, in the case of small molecular graphs, the sum function can capture representative elements in the graph (as well as the max function) and can learn to store information of subgraphs of different sizes (as well as the concatenation function), if these strategies are relevant for DTI prediction. Regarding the hierarchical pooling, it seems that standard GNN can also leverage information of substructures of different sizes in a hierarchical manner. Indeed, the iteration of the nodes update can already provide neighbouring information to each node in a hierarchical manner.

### 6.3 Evaluation of protein sequence encoding architectures

Up to now, we only explored the modification of the GNN architecture encoding molecular graphs, but did not observed significant improvement in the prediction performance of the resulting chemogenomic network. However, the performance bottleneck could be related to the protein sequence encoder rather than the molecular graph encoder.

Therefore, we replaced the stacked CNN block that encodes the proteins in the standard chemogenomic network by a biLSTM layer on top of a convolution layer. Indeed, the bi-LSTM can locally integrate the detected patterns by the convolutional layer to provide local contextual information, both forward and backwards, whereas stacked convolutional layers process protein sequences by detecting local patterns, patterns of patterns and so on.

We varied the dimension of the amino acids learnt representations ({10, 50, 100, 500}) and the amount of dropout ({0., 0.1, 0.3, 0.6}), to search for the best performance.

However, the best performance reached by this architecture was identical that obtained while using stacked convolutional layers of the standard chemogenomic network, as shown in Table 5 at the row named “CN-biLSTM”.

At this stage, we conclude that encoders architecture refinements, both for molecules and proteins do not appear as promising directions to enhance DTI prediction with the chemogenomic framework.

### 6.4 Evaluation of the neuron architecture combining the learnt representations of molecules and proteins

We now propose to investigate the use of an attention mechanism in the CNN protein sequence encoder, which modifies the simple concatenation used as *Comb* operation in the standard chemogenomic network.

We first implemented the attention mechanism proposed by Tsubaki et al [78] and displayed in eq. 3, which resulted in a significant decrease of performance for the four *S*_1_, *S*_2_, *S*_3_, and *S*_4_ settings. This might be due to the fact that this attention mechanism is highly non-convex, since it relies on the dot product of the two learnt representations, resulting in a rugged optimisation landscape.

Therefore, instead of using an attention mechanism calculated only for the amino acids representations, we turned to an attention mechanism that computes the pairwise representation **h**_*pair*_ based on the concatenation [**h**_*prot*_, **h**_*mol*_] of the protein **h**_*prot*_ and molecule **h**_*mol*_ learnt representations, following:

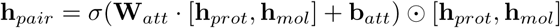

In the previous equation, *σ* refers to the sigmoid function and ⊙ to the element-wise multiplication. This self-attention mechanism differs from that of Tsubaki et al. in two ways. First, it selects learnt abstract features at the protein and molecule level, whereas the precedent attention mechanism learnt attention weights on abstract features at the amino acid level. Second, this mechanism operates on the concatenation of the protein and molecule abstract features, and jointly selects the most important protein and molecule features with respect to the prediction task. This appears rather intuitive since the protein-ligand interaction mechanism relies on both partners. However, in the present case, the selected features are abstract, and therefore, not easily interpretable.

As reported in Table 5 (at the row named “CN-pairwiseAtt”), this did not result in performance improvement, meaning that such soft attention mechanism does not provide better representation power. It is most likely that the self-attention mask “*σ*(**W**_*att*_ · [**h**_*prot*_, **h**_*mol*_] + **b**_*att*_)” produces a uniform attention weight distribution in most cases.

Up to now, we only explored modifications of the neuron network architecture to encode (protein, molecule) pairs from their raw representation. We conclude that searching for more sophisticated architectures is not the right direction to improve the prediction performance of the the chemogenomic neuron network, at least as observed on our DrugBank-based benchmark dataset.

In other words, since the performance does not see to be limited by the representation power of the chemogenomic neuron network architecture, the bottleneck seems to be related to data featurisation, diversity and quantity.

In particular, a feed forward neuron network on top of expert-based features is highly competitive. Therefore, we evaluate the simultaneous use of expert-based and learnt features in the next section.

### 6.5 Evaluation of the combination of expert-based and learnt features

In the first experiment of this Results section, we showed that the reference FNN trained with expert-driven features as inputs, outperforms the standard chemogenomic neuron network in some settings. Therefore, we tested an architecture that intend to leverage information from these two forms of inputs using appropriate neuron network components: (i) the expert-based features and (ii) the raw data (molecular graph and protein sequence). The idea is that such a network may benefit from the best of both forms of representations, and assess the degree with which each representation is efficient to predict DTIs. Therefore, it is expected to perform better than the two types of representations (i.e. expert-based and learnt) taken separately.

In the following discussion, we considered the same expert-based features that were used as input of the FNN-based model in the first experiment.

Furthermore, there are several possibilities of neuron network architectures to integrate multiple views of proteins and molecules, in particular, expert-based and learnt features, into a final pairwise representation. We considered the two architectures displayed in Fig 4.

**Figure 4:**
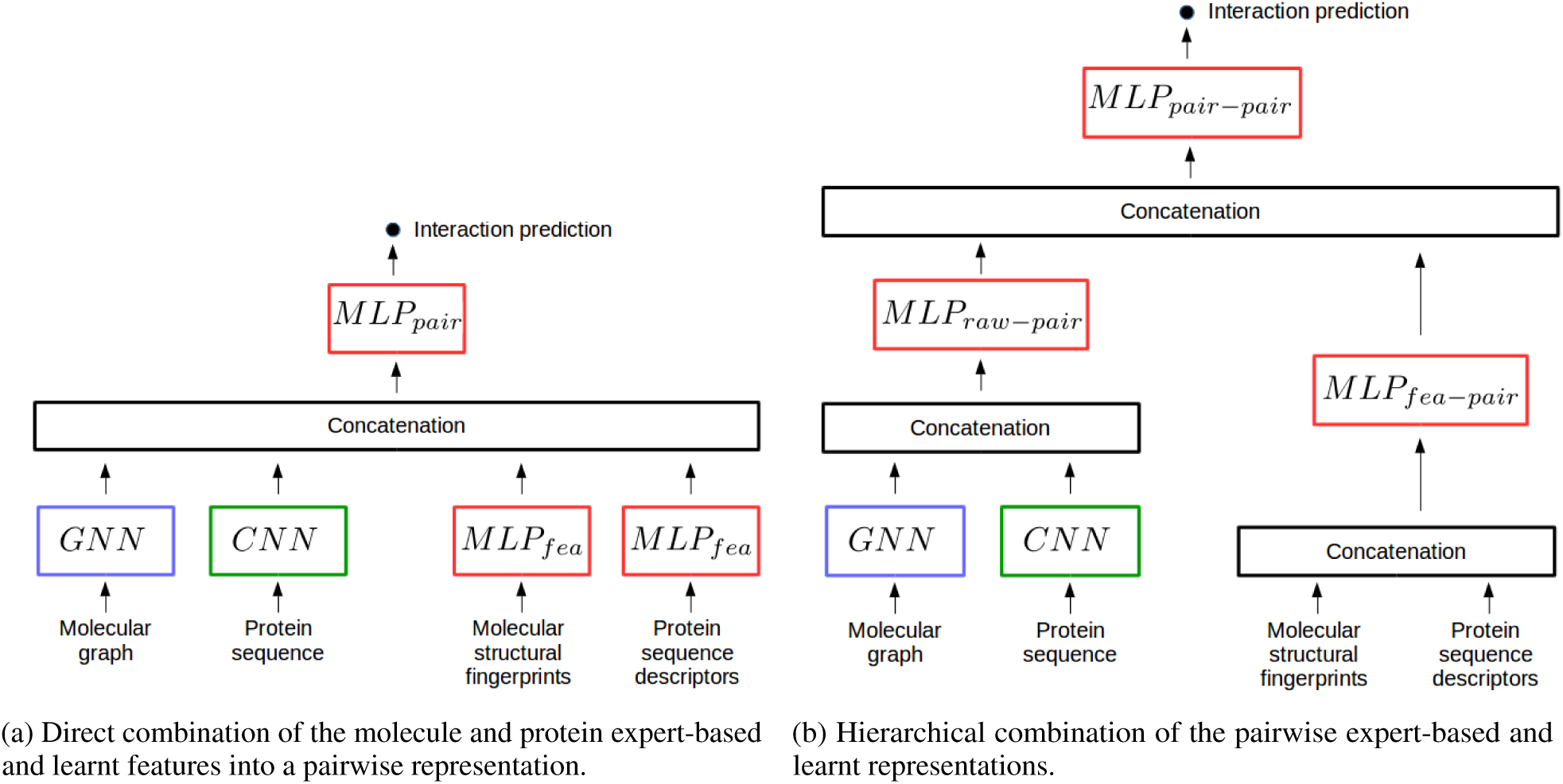
Two types of chemogenomic neuron networks combining expert-based and learnt features. (a) Direct combination of the molecule and protein expert-based and learnt features into a pairwise representation. (b) Hierarchical combination of the pairwise expert-based and learnt representations.

The architecture in Figure 4a combines in one step the four types of data representations: (i) the molecular graph, (ii) the expert-based molecule features, (iii) the protein sequence and (iv) the expert-based protein features. Regarding the multi-layer perceptron which processes molecule and protein expert-based features (*MLP*_*fea*_ in Figure. 4a) and that processing the concatenation of the four representations (*MLP*_*pair*_ in Figure. 4a), we explored the hyper-parameter spaces by varying the number of layers (from 1 to 3), the number of hidden units (in {10,100,200,1000,2000}) while promoting pyramidal architectures (as motivated in [70]) and the quantity of dropout (in {0,0.3,0.6,0.9}).

To rule out the possibility of an implementation error, we performed two sanity checks. First, we checked that we recover the performance obtained by an MLP with expert-based features when we set to one the dimension of the expert-based features (i.e. we silent the protein sequence and molecular graph encoders while keeping the same implementation). Similarly, we also checked that we recover the performance obtained with expert-based features while silencing *MLP*_*fea*_ and *MLP*_*pair*_ in the same manner.

The performance reached by such a model is significantly lower than that obtained individually by the chemogenomic neuron network and an FNN with expert-based features, on the four *S*_1_, *S*_2_, *S*_3_, *S*_4_ settings of the *DBEC* dataset (data not shown). A possible explanation is that simultaneous training of the *MLP*_*fea*_ and *MLP*_*pair*_ networks with molecular graph and protein sequence encoders is an ill-conditioned non-convex optimisation problem.

To address this issue, we first separately pre-trained the pairs of neuron modules encoding expert-based and learnt features, before re-training them together on the same training data. More precisely, with respect to the scheme represented in Figure 4a, the *MLP*_*pair*_ network is trained with the GNN and CNN networks alone, then with the *MLP*_*fea*_ for proteins and molecules alone. The pre-trained GNN and CNN, and *MLP*_*fea*_ networks are finally retrained together, as in the architecture displayed in Figure 4a. The parameter starting values of protein and molecule networks are those optimised in the pre-training phase, whereas the starting values of parameters in the *MLP*_*pair*_ are randomly initialised. With this procedure, we could only retrieve the performance reached when considering the expert-based representation alone, even when increasing the width and depth of *MLP*_*pair*_.

To further explore joint optimisation of neuron modules for expert-based and learnt feature extraction, we considered the architecture in Figure 4b. It combines the pairwise expert-based and learnt representations such that both representations are combined only through a single relatively small layer. Typically, *MLP*_*pair*-*pair*_ (cf. Figure 4b) is composed of a single hidden layer of 50 hidden neurons. In the following, we call this model the *CN* -*feaMLP* approach.

We varied the neuron architecture and dropout probability of *MLP*_*fea*-*pair*_ similarly as for *MLP*_*fea*_, and the ones of *MLP*_*raw*-*pair*_ and *MLP*_*pair*-*pair*_ similarly as for *MLP*_*pair*_, in the architecture displayed in Fig. 4a.

The performance of such a network are reported in Table 5 at the row named *CN* -*feaMLP*. Remarkably, this approach significantly outperformed the standard chemogenomic neuron network for the *S*_1_ setting, while performing similarly on the three others. More precisely, the best-reported performance on the *S*_1_ setting is the one reached by the reference FNN that simply uses expert-based features as input.

To further challenge this approach, we assessed the performance on the four *S*_1_, *S*_2_, *S*_3_ and *S*_4_ settings using a 5-fold nested cross-validation scheme, with a test and train “positive:negative” ratio of 1:5. The performance on *DBEC* and *DBH* are reported in Figure 5 next those reached by the reference methods and the standard chemogenomic network of the present study (CN). For the *DBEC* dataset, the *CN* -*feaMLP* model outperforms both the standard chemogenomic neuron network and the FNN reference model (with expert-based protein and molecule descriptors as inputs) on *S*_1_ and *S*_2_, and performs similarly, yet slightly better, for the *S*_3_ and *S*_4_. However, the performance is still significantly lower than that of *NRLMF*, on this small dataset. Therefore, we conclude that the *NRLMF* approach should be used on relative small datasets.

**Figure 5:**
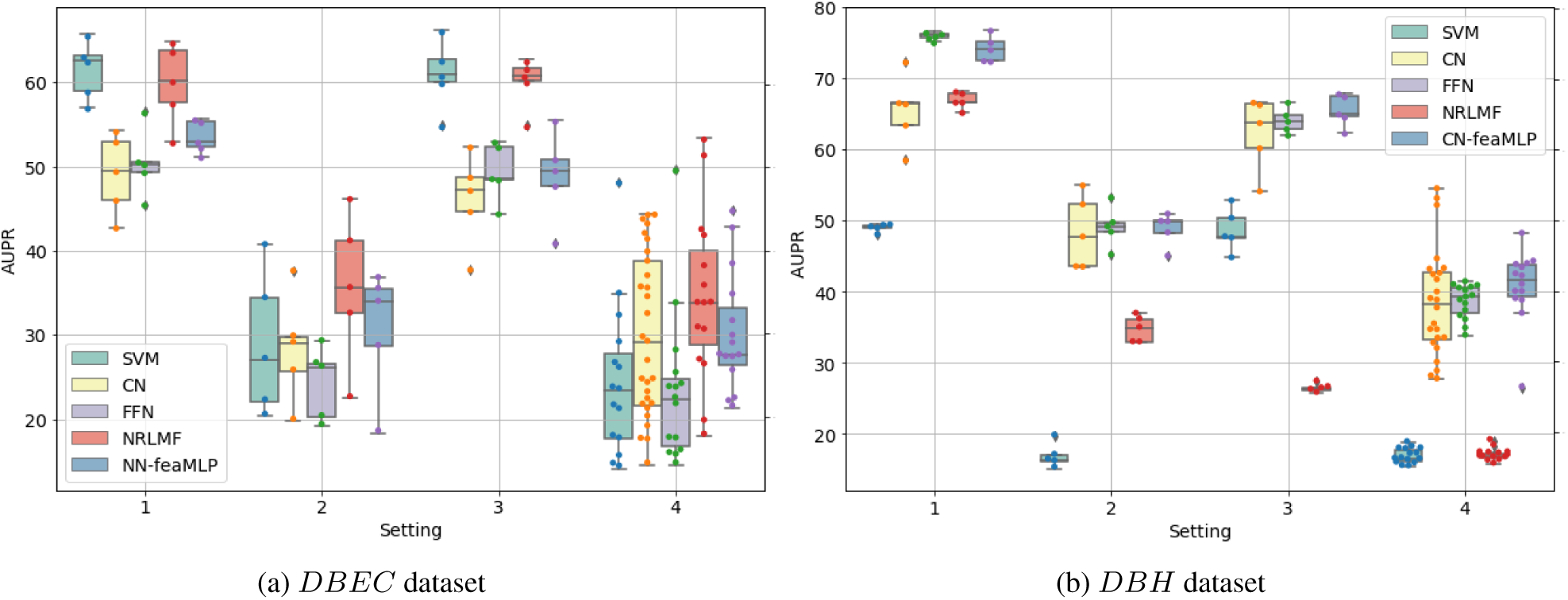
AUPR scores obtained via 5-fold nested cross-validation for the four *S*1, *S*2, *S*3, and *S*4 settings. The performance of *CNN* -*feaMLP* are compared to the reference shallow methods and to FNN and CN as reference methods for deep learning. For each setting, the order from left to right in which the results of the different methods are displayed corresponds to that of the legend in the corner of the figures.

However, on the larger *DBH* dataset, the *CN* -*feaMLP* model outperforms the standard chemogenomic neuron network model for the *S*_1_, but it does not overtake the FNN approach. For *S*_2_, *S*_3_ and *S*_4_, the *CN* -*feaMLP* approach reaches slightly better performance than both the standard chemogenomic neuron network and the reference FNN method.

Overall, we conclude that, for relatively large datasets like *DBH* (that are widely available our-days), the *CN* -*feaMLP* model provides the best DTI prediction performance in chemogenomic studies. Moreover, we believe that integrating other multiple description views of molecules and proteins in the same model, is an interesting domain for future developments.

Within the same type of idea, in the next subsection, we explore another relevant “data-oriented” direction to enhance the performance of the chemogenomic neuron network: transfer learning for implicit data augmentation. Indeed, transfer learning is a way to combine different tasks related to the same types of data (proteins or molecules in our case), in the aim of providing additional information about the data, based on the information they bear for different prediction tasks.

### 6.6 Evaluation of transfer learning for chemogenomics

The principle of transfer learning is to gain knowledge while solving one problem and applying this knowledge to a different but related problem. In fact, prediction of DTIs in the context of chemogenomics can already be viewed as a form of transfer learning. Indeed, we can formulate the problem of predicting DTIs at a large scale in the protein and molecule spaces as a problem of predicting ligands for each protein, while learning on all the problems of predicting ligands for all the other proteins. In this particular case, all the tasks are formally identical, since they correspond to predicting ligands for proteins.

In the present section, we investigate the efficiency of transfer learning in the context of chemogenomics, using tasks that are not related to the prediction of ligands for proteins within the same dataset, but other tasks involving proteins and expected to implicitly contain valuable biological information for solving the chemogenomic problem.

In the context of deep learning, transfer learning can take two forms: pre-training, or co-training encoders on separated larger prediction tasks (even unsupervised ones). Leveraging the information in larger datasets is aimed to enhance performances on the source task.

Co-training refers to the parameter sharing strategy in which we train several prediction tasks simultaneously, while task-specific neuron encoders share information in order to enable them to learn a richer set of features.

We tested co-training on two prediction tasks which are directly related: one based on the “SCOPe” database [95] on which we classified 13.963 proteins over seven types of 3D fold (alpha proteins, beta proteins, alpha and beta proteins, multi-domain proteins, membrane and cell surface proteins and peptides, small proteins), and one a “secondary structure” dataset [67] composed of 4388 proteins in which all amino acids are assigned to one of the eight possible types of secondary structures. Thus, these two prediction tasks are explicitly related since they both characterise protein folding, but at different level.

The co-training of the protein sequence encoder on the SCOPe and “secondary structure” datasets did not yield to an improvement of performance (data not shown), although these two tasks are strongly related. A similar observation was made in the Natural Language Processing field [96]. Co-training learnable feature encoders is likely to be too restrictive and therefore, is probably efficient only in few specific cases. Therefore, we did not further consider co-training in this study.

Pre-training, also called “curriculum learning”, is a procedure in which the model is first trained on a large dataset (called the source dataset) to learn a rich set of features that are optimised to make predictions for source task trained on a large dataset. A sketch of curriculum learning in the case of the chemogenomic network can be found in Fig. 6.

**Figure 6:**
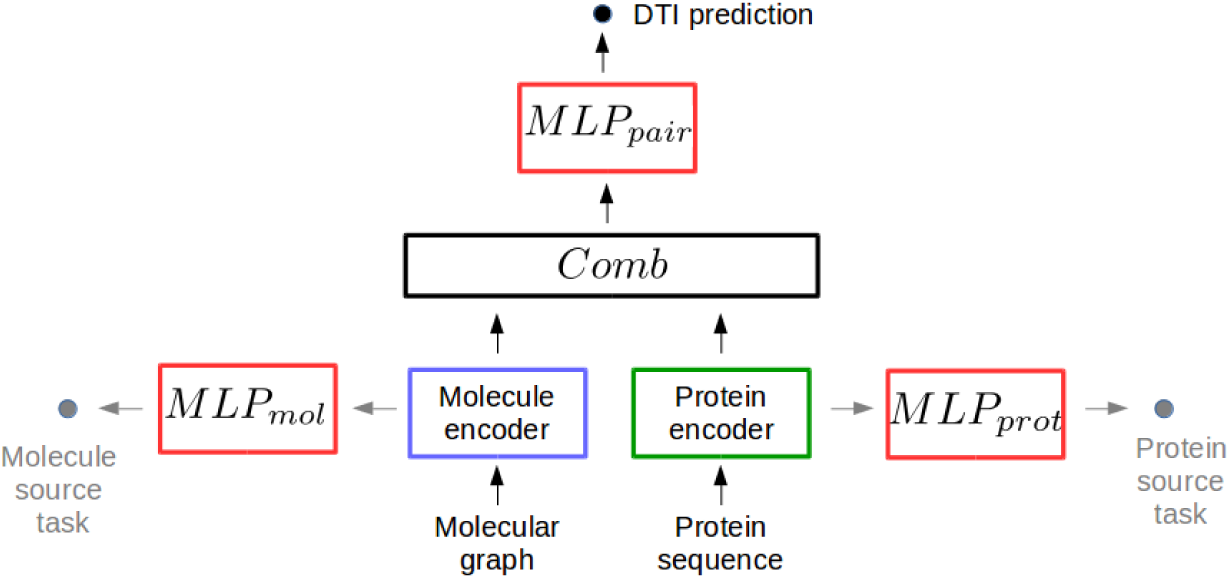
Sketch of curriculum learning with the chemogenomic neuron network. First, the learnable protein sequence and molecular graph encoders are optimised with molecule- and protein-specific source tasks (grey path). In a second phase, the pre-trained sequence and molecular graph encoders are used (either “frozen” or re-trained, to fit the chemogenomic prediction task (black path).

Next, this pre-trained model is fine-tuned by re-training the weights on the smaller dataset of interest (named the target dataset). This means that, when training for the target task, the initial parameters of the encoder are not randomly chosen, but are assigned to their optimised values for the source task. In such a procedure, the model fits the large input data available for the source task, and fine-tunes its weights in a second phase to adapt to the smaller dataset of the target task.

Alternatively, it is also possible to “freeze” (i.e. not retrain) the pre-trained molecule and protein encoders, and train only the final FNN that take as input the representations learnt from the pre-trained encoders. In the case where the source and target datasets are related “enough”, this could improve predictions on the target dataset.

For instance, Paul et al [97] performed such transfer learning in a chemoinformatics problem related to organic solar cell screening (only chemicals are screened).

Before discussing our results, we start with general observations about implementation of curriculum learning with the chemogenomic neuron network.

First, we performed a sanity check to challenge our implementation of curriculum learning. More precisely, we checked that a pre-trained model reaches immediately best performance when re-using the pre-trained network on the same train and test data, with or without freezing all the layers.

Second, we found that curriculum learning made often the training much shorter, in terms of the number of training epochs in the second phase. Indeed, pre-trained encoders are expected to be closer to the optimal solution than encoders whose weights are randomly initialised.

Third, we considered to pre-train and load per block the three sub-networks of the chemogenomic neuron network: (i) the protein sequence encoder, (ii) the molecular graph encoder, and (iii) the prediction sub-network, i.e. the stacked fully connected layers of the FNN that outputs the DTI predictions. Furthermore, we varied the number of layers and neurons in the prediction sub-network in order to search for the best architecture when freezing one or the two encoders. Indeed, an MLP may need a relatively large width and depth to leverage the representations extracted by pre-trained frozen encoders.

Finally, it is of general interest to explore whether transfer learning can improve the prediction performance of the standard chemogenomic network. However, we chose to focus on the *DBEC* dataset. Indeed, it seemed more critical to perform data augmentation on a small dataset like *DBEC*, and because we already showed that deep learning methods could outperform reference shallow machine-learning methods (SVM or *NRLMF*) on larger datasets like *DBH*. Therefore, the goal was to evaluate whether transfer learning improves prediction performance of the chemogenomic neuron network on *DBEC*, and whether these improvements allowed this deep learning method to reach the performance of reference shallow methods.

In a first attempt, we considered to pre-train the three parts of the chemogenomic neuron network (the protein sequence encoder, the molecular graph encoder and the prediction sub-network) on the larger *DBH* dataset in order to use it for prediction on the smaller *DBEC* dataset. This setting is slightly different from that shown in Fig. 6: here, a single source task is used to pre-train the whole CN network (protein encoder, molecule encoder and FNN), corresponding to predicting DTIs on the larger *DBH* dataset, and use the optimised parameters as starting values (instead of random values) to re-train the whole CN on the smaller *DBEC* dataset. This led to a loss of performance on *S*_1_ (0.2 of AUPR instead of 0.4).

Interestingly, we found that this loss of performance arouse from the protein encoder pre-trained on *DBH*. Indeed, pre-training on *DBH* and freezing the protein sequence encoder, while training the molecular graph encoder and prediction sub-network directly on the target task with randomly initialised parameters, led to the same decrease in AUPR score. On the contrary, when pre-training and freezing the molecular graph encoder while training the protein sequence encoder and the prediction sub-network directly on the target task, we obtained similar or better performance than with direct training of the chemogenomic neuron network, as shown in Table 5 at the row named “curriculum learning”. In particular, the performance obtained on *S*_1_ and *S*_3_ were significantly better. This means that DTI prediction with the chemogenomic neuron network may benefit from a molecular graph encoder pre-trained on larger datasets.

The two prediction tasks (*DBH* and *DBEC*) are formally identical, but their training data have different characteristics. Indeed, *DBEC* contains E. coli proteins and their ligands, whereas *DBH* contains human proteins and their ligands. Therefore, the two datasets may present critical statistical bias in protein sequences characteristics, since the distribution of protein sequences in bacteria is expected to differ from that of human proteins. However, both datasets contain small organic drug-like molecules. These remarks may explain why a loss of performance is observed when pre-training with human proteins, while pre-training only the molecular graph encoder on the *DBH* dataset is suitable to predict DTIs on the *DBEC* dataset.

When we pre-trained the encoders without freezing the parameters in the second phase, and use an initial learning rate in the second phase one order of magnitude lower than the one used to train directly the chemogenomic neuron network (i.e. initial learning rate set to 10^*-*4^ instead of 10^*-*3^), the performance was lower than that obtained when training the chemogenomic neuron network from scratch. In particular, the average AUPR is 0.3 instead of 0.4 on *S*_1_. This means that encoders’ weights pre-trained on *DBH*, in particular those of the protein encoder, are not suitable starting points for fine-tuning on *DBEC*.

To investigate these conclusions, we selected larger and chemical and protein datasets to pre-train the encoders with source tasks that are now different in nature, although related to the target task.

The PCBA dataset is an excellent choice for pre-training a graph neuron network encoder for molecules since it contains the information about a hundred of bio-activities for hundreds of thousands of molecules. This source task for molecules is to predict whether they are active or not. To our knowledge, there is no such evident choice for protein datasets. We considered the sub-cellular localisation dataset [67], containing six thousand proteins labelled with one of the twelve sub-cellular localisation, called the *CellLoc* dataset in the following (see Materials and Methods for a description of these datasets). This source task for proteins is to predict their cell localisation. Recall that the target task is to predict DTIs in the *DBEC* dataset.

We considered pre-training only two blocks of the chemogenomic neuron network, (i) the protein sequence encoder and (ii) the molecular graph encoder in order to use them for DTI prediction on *DBEC*. When pre-training the molecular graph and protein sequence encoders on the *PCBA* and *CellLoc* datasets respectively, and freezing them before fitting the FNN model for DTI prediction on the *DBEC* dataset, we recovered the performance reached by direct training of the standard chemogenomic neuron network on *DBEC*. More precisely, we recovered a performance of 0.4 in AUPR only with a prediction neuron sub-network with three hidden layers of 1000, 500 and 100 hidden neurons each. However, we obtained 0.28 of AUPR with a prediction neuron sub-network made of a single hidden layer of 100 hidden units. This means that the prediction sub-network needs to be complex enough to leverage the information in the representations learnt by the pre-trained encoders. In other words, these pre-trained encoders are informative with respect to the *DBEC* prediction task, but not as suited as encoders trained on the *DBEC* prediction task itself.

To study curriculum learning in more details with the chemogenomic neuron network, we evaluated pre-training of the molecular graph encoder on *DBH* and *PCBA* via 5-fold nested cross-validation with a test and train “positive:negative” ratio of 1:5 on the four *S*_1_, *S*_2_, *S*_3_ and *S*_4_ settings of the *DBEC* dataset. In the case of pre-traning with *DBH*, the strategy is slightly different from that presented in Figure 6. In fact, the standard chemogenomic network is pre-trained with *DBH*, but only the parameters of the molecule encoder are retrained on *DBEC* (i.e. parameters of the protein encoder and of the FNN are randomly initialised). In the case of pre-training with *PCBA*, since this dataset only contain bio-activities for molecules, the pre-training step only involves the molecule encoder and the FNN, and the source task is to predict the PubChem assay bio-activities. Then, the parameters of the molecule encoder are kept for retraining the chemogenomic network on *DBEC*.

The performance are reported in Figure 7 with those of the reference methods. The final model obtained when pre-training the molecular graph encoder on the *PCBA* (resp. *DBH*) dataset is called *CN* -*currPCBA* (resp. *CN* -*currDBH*) in Fig.7

**Figure 7:**
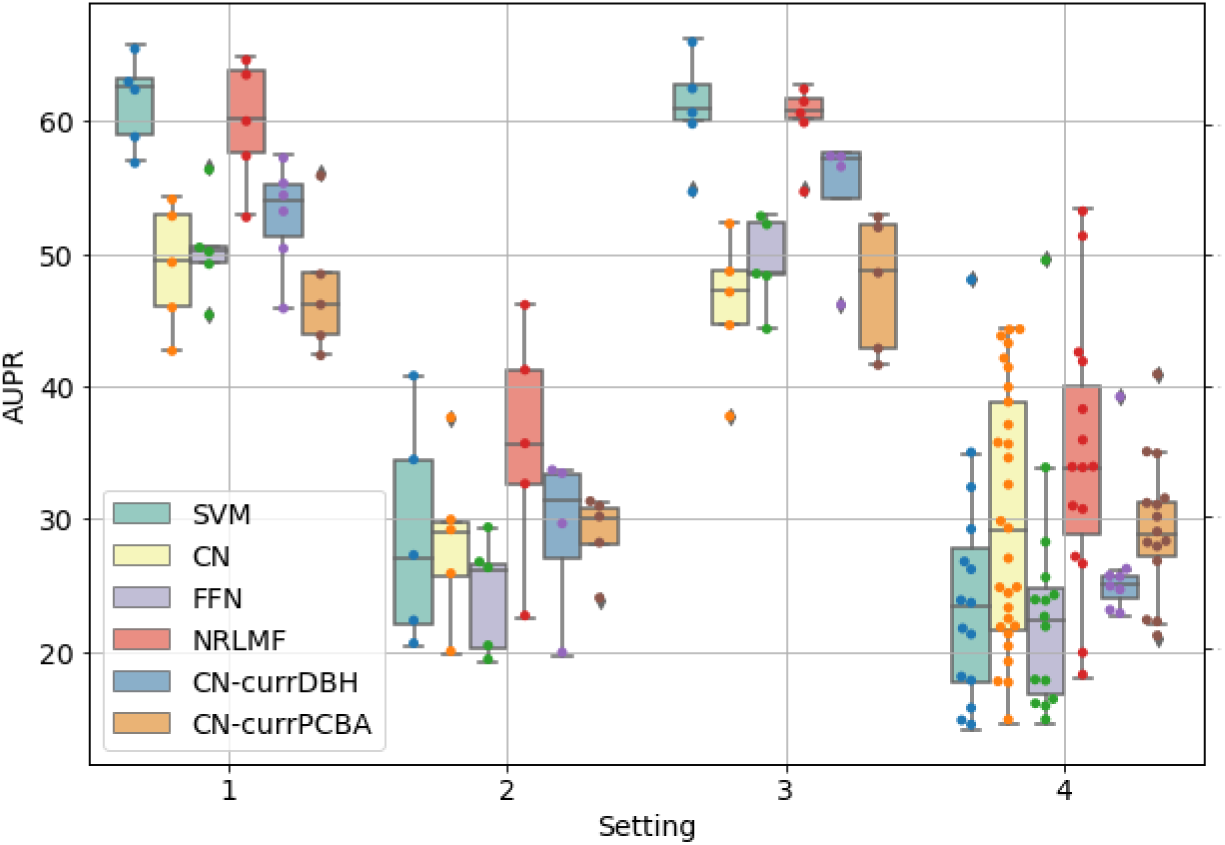
AUPR scores obtained with a 5-fold nested cross-validation scheme for the *S*_1_, *S*_2_, *S*_3_, and *S*_4_ settings on the *DBEC* dataset. The performance of *CNN* -*currPCBA* and *CNN* -*currDBH* are compared to the reference shallow methods (SVM and *NRLMF*) and to FNN and CN as reference methods for deep learning. For each setting, the order from left to right in which the results of the different methods are displayed corresponds to that of the legend in the corner of the figures.

Pre-training the molecular graph encoder on *PCBA* yielded to similar performance than direct training on *DBEC* for the four settings. We thus confirmed with 5-fold nested cross validation the observations previously made on a single train/validation/test split for the four settings.

Interestingly, pre-training the molecular graph encoder on *DBH* resulted in a better model than that trained directly on the *S*_1_, *S*_2_ and *S*_3_ of the *DBEC* dataset (CN and *CN* -*currDBH* in Figure 7), as observed in the previous experiment with a single train/validation/test split. Although the proteins involved in *DBH* and *DBEC* follow different statistics, the global prediction tasks are indeed strongly related, so that transfer learning can be observed between the two tasks.

Future studies could consider larger source datasets. For instance, we could consider an unsupervised learning task as a source task such as a reconstruction task like in auto-encoders [37, 98] (predicting the original input from the hidden representation), or an adversarial task [99] (distinguishing real data from fake data, such as non-valid molecules or randomly generated protein sequences).

However, fig. 7 shows that, in general, the reference methods SVM or *NRLMF* still led to the best performance on this small *DBEC* dataset, and even with transfer learning, deep learning methods did not outperform these reference machine-learning methods.

## 7 Conclusions

In the present work, we investigated deep learning methods for chemogenomics. In particular, we studied representation learning methods for molecules and proteins based on deep learning networks in the context of drug-target interaction prediction. To this end, we first defined the chemogenomic neuron network which consists of a feed-forward neuron network (FNN) taking as input the combination of molecular and protein learnt representations based on molecular graph and protein sequence encoders. More precisely, data representation are learnt from the raw description of the considered data, in order to maximise prediction performance.

For large datasets, we show that this standard chemogenomic neuron network (with GCN as graph molecule encodre and CNN as protein sequence encoder) outperforms state-of-the art reference machine-learning methods not using deep learning, and is competitive with the reference FNN models using expert-driven descriptors as inputs, particularly in orphan settings like *S*_2_, *S*_3_ and *S*_4_, for which we found that the chemogenomic neuron network reached the best performance.

Moreover, we investigated the latest developments in graph representation learning, and point out critical issues that question the efficiency of more complex methods to learn representations in the context of chemogenomics. Although the diversity of neuron network architectures covered in the present work, and their applications in virtual screening, are not exhaustive, we can still conclude that the current research in the neuron network architecture for encoding molecular graphs and protein sequences does not seem a promising direction to improve drug virtual screening in the chemogenomic framework. Indeed, we observed that promising modifications of the graph encoder (inspired from other fields of graph learning) to learn molecule descriptors, and of the protein sequence encoder and pairwise representation encoder, as parts of the chemogenomic neuron network, did not yield to significant improvements in prediction performance, on our DrugBank-based dataset. Nevertheless, GNN is a very dynamic field and new developments might lead to improvements in chemogenomics in the future [100].

Based on our results, integration of heterogeneous data sources appeared as a more encouraging direction to follow. We explored whether integration of heterogeneous data sources could improve the performance of the standard chemogenomic network. We investigated two directions for integration of different data sources: (1) the combination of learnt and expert-driven features in a single neuron network, and (2) transfer learning with larger source prediction tasks. To our knowledge, the present work is the first to consider and evaluate data augmentation techniques in chemogenomics with the representation learning-based approaches.

Integrating learnt and expert-driven features in the same network yielded to substantial improvements. Therefore, for future directions, we recommend to assign efforts in the design of neuron network that integrate multiple views of molecules and proteins in the same model. However, in the case of small datasets like *DBEC*, these performance improvements did not allow to reach those of reference shallow machine-learning methods (SVM or *NRLMF*).

In contrast, we report that curriculum learning did not result in strong performance improvement on the small *DBEC* dataset, meaning that a promising direction to improve the chemogenomic neuron network is at first to integrate various data views, rather than to increase the amount of auxiliary data.

Our study builds on and provides new insights on the chemogenomic neuron network initially proposed by several authors [101, 78, 76]. In particular, these studies did not justify their choice of neuron network architecture via the comparison of prediction performance of various architectures, did not consider most recent developments in graph neuron networks, and did not compare to established state-of-the-art machine-learning methods (SVM or *NRLMF* for shallow methods, and FNN with expert-based descriptors for deep learning methods).

In addition, we used different settings (*S*_1_, *S*_2_, *S*_3_, *S*_4_) in order to better assess the performance of the considered methods in a panel of scenarios that are encountered in practice. Indeed, real case studies usually search proteome-wide off-targets for new drugs or drugs for new therapeutic targets. They are closer to the orphan settings *S*_2_, *S*_3_, or *S*_4_ that to the *S*1 setting. Therefore, measuring prediction performance only on datasets similar to *S*_1_ overestimates the performance that can be expected in real case studies [28].

We emphasise that FNN taking as input standard expert-based molecular fingerprints and protein sequence descriptors is a good default method for prediction of DTI on relatively large datasets like *DBH*, and therefore, should be considered as a reference simple method that leads to state-of-the-art performance, and to which future chemoinformatic studies should be compared. Indeed, we reported that feed-forward neuron networks applied to expert-based descriptors are competitive and even yield to the best performance in the raw setting (*S*_1_) of proteome-wide DTI prediction. Moreover, remember that we did not fine tune this model, and most importantly, we did not considered the most relevant expert-crafted descriptors for the DTI prediction task which leaves some space for improvement and reinforces the interest of this approach. Indeed, combining multiple types of fingerprint descriptors provides the best performance [102, 90, 103, 104] and that the best fingerprints to retrieve a protein-ligand interaction depends on the protein target [102, 90]. Regarding protein descriptors, proteochemometric attributes [105] may be more suitable for DTI prediction.

Besides, neuron network-based models were not efficient on the small dataset *DBEC* on which shallow methods performed the best. In particular, *NRLMF* appeared to be the method of choice in this case. Nevertheless, we think that most models should be evaluated on relatively large datasets, with thousands of recorded interactions, since the publicly available databases now reach this size. In particular, evaluating new methodologies on small datasets scrambled the examination of new methodologies, like in [78]. In practical applications of DTI prediction, we finally suggest to cconsider an ensemble learning strategy, combining the predictions of *NRLMF*, FNN and the chemogenomic neuron network, since they individually perform the best on different types of the problems.

Most importantly, we conclude from our observations that a promising future research direction is to integrate heterogeneous sources of data such as various bioactivity datasets, and independently, multiple molecule and protein attribute views. Indeed, all data sources are expected to be noisy (missing data, experimental noise) but with different noise pattern, and to cover different sub-parts of the chemical space. Therefore, combining data sources may have a regularisation effect as well as provide information from a wider sub-part of the chemical space.

## Acknowledgement

The authors would like to thank the CBIO lab members, in particular Chloé-Agathe Azencott, for many valuable discussions.

This work was supported by the Mines ParisTech, and by Ministère de l’Industrie.

